# Spatiotemporally selective ATP events from astrocytes encode injury information and guide sustained microglial response

**DOI:** 10.1101/2022.06.21.497103

**Authors:** Yue Chen, Pengwei Luan, Juan Liu, Yelan Wei, Zhaofa Wu, Miao Jing

## Abstract

Brain injuries, either directly result from external assaults or are accompanied with diseases, initiate a cascade of intrinsic responses for damage management, with resident microglia as one of the key responders in the early phase of reaction. Changes in microglia including their motility and activities can be tuned according to injury intensity and position, indicating such injury information is precisely encoded and actively presented. Also, microglia in a broad area can perform sustained migration towards a transient and local injury, suggesting a signal amplification process may exist that bridge differences between injury and microglial migration in time and space. Currently, the molecular identity and underlying mechanism for injury encoding, amplification and presentation have not been fully elucidated yet, although candidate molecules like ATP are linked with both injury and microglial response. Based upon our recent technique advancement in engineering novel genetically-encoded ATP sensors, we here identified and characterized that in the cortex of awake mice in vivo, a new type of spatiotemporally selective ATP events, referred to as Inflares, was selectively evoked after brain injuries, which were actively and repeatedly generated from astrocytes in a Ca^2+^-dependent manner through the opening of pannexin 1 channel. Functionally, Inflares amplified local injury over time and space with their persistence and widespread distribution, and provided continuous directionality that was necessary for guiding microglial migration. Excessive Inflares in pathological injuries drove microglial dysfunction and caused secondary damage, whereas blocking Inflares successfully reversed pathological changes and benefited the outcome of ischemic stroke. Together, we identified the internal mechanism that encoded and presented injury information, and provided rational target for treating injury-related diseases.

## Introduction

In all living organisms, the ability to precisely sense and respond to injury, especially an injury in the central nervous system (CNS), is critical for survival. Injuries caused by external traumatic damage, e.g. from traffic accidents, and the internal perturbations like reduced oxygen and blood supply during ischemic stroke, continue to be the leading cause of death worldwide, and with high rate of disability that place huge burden on society and public health (Donkor, 2018; Taylor et al., 2017). Meanwhile, CNS-relevant diseases like neurodegenerative diseases, brain tumor that generate aggregation of toxic products, physical stress or competition in energy supply, all lead to damage and death of healthy cells and lead to clinical symptoms (Mattson, 2000; Perry et al., 2016; Stylianopoulos et al., 2012). Different from being passively damaged, brains can initiate an active response in coping with damage and promote tissue recovery, which is organized in a cascade of cellular events that are achieved by cooperation between cells (Gadani et al., 2015). Such active response to injury is under precise regulation, and abnormal injury response may lead to failure in controlling the damage and subsequently affect disease outcome. In theory, an appropriate response to injury requires high sensitivity and precision to locate even a minor damage at cellular or sub-cellular level, while also amplifying the response signal over time and space to recruit enough responder cells for effective management. In addition, such signal detection and transmission should be able to maintain robustness in the dynamic environment *in vivo*. Yet, how these features are implemented is largely unclear.

Microglia as the brain-resident immune cells are the principal mediators of the injury response. Besides surveying the parenchyma with low motility at resting states, microglia can quickly extend processes and exhibit directional migration to local sites of injury, followed with proliferation and activity changes in mediating diverse injury response, including but not limited to phagocytosis of cell debris, initiation of inflammatory response and secretion of growth factors for tissue repairing (Bianco et al., 2005; Carbonell et al., 2005; Davalos et al., 2005; Kreutzberg, 1996; Nimmerjahn et al., 2005; Shieh et al., 2014; Walz et al., 1993). Interestingly, damage on local blood barriers, or precise introducing sub-cellular level of injuries, are able to mobilize a broad region of microglia to the site, even those hundreds of micrometers away from the damage (Ahn et al., 2018; Carbonell et al., 2005). Meanwhile, after the removal of initial damage, microglia could still keep migration for hours or even days (Katayama et al., 2012; Miller et al., 2019). These indicate that the initial damage is amplified in time and space, which transferred into a sustained signal that keep guiding microglial response. Such process may ensure the effective response to damage and prevent its exacerbation, yet the underlying mechanism is still unclear.

Pioneer work uncovered that microglia migration critically depends on the purinergic signals ATP, with the degradation of extracellular ATP, or blocking/knockout microglial receptor P2Y12 prevent their migration to the injury site (Duan et al., 2009; Haynes et al., 2006; Honda et al., 2001; Miller and Stella, 2009; Wu et al., 2007). In cultured microglia, local application of ATP could mobilize and recruit microglia, mimicking the response after injury in vivo (Dou et al., 2012; Honda et al., 2001). Besides activating microglia, ATP has also been linked with injury. Early studies using microdialysis of brain cerebrospinal fluid sample, as well as biochemical analysis of ATP level in homogenate brain tissue, have reported the increase of ATP level follows brain injury (Bell et al., 1998; Sullivan et al., 1998), establishing the correlation between injury and ATP. Using bioluminescence reporter for ATP, it has been shown that a bulk increase of ATP exists surrounding the peri-lesion site (Samuels et al., 2010; Wang et al., 2004).

Further, ATP have been reported to be released from damaged apoptotic cells follows the concentration gradient (Chekeni et al., 2010; Elliott et al., 2009; Gribble et al., 2000), which could act as damage-associated molecular pattern (DAMP) to active microglia for phagocytosis and release of inflammatory mediators (Bianco et al., 2005; COUILLIN et al., 2013). Together with the fact that brain injury resulted in local cell apoptosis, and the ability of artificially generated ATP gradient in recruiting microglia in vitro, current theory viewed ATP as the “find me” signal from apoptotic cells (Ravichandran, 2010), and passively diffuse away to recruit microglia as responders. This simplified view, however, cannot reconcile with other evidence of the microglia response and brain ATP regulation. First, enzymes that degrade ATP, including CD39 and CD73 are highly expressed on the membrane of multiple cell types in the brain (Zimmermann, 1996), thus limit the effective time and diffusion area of ATP after release. Second, some minor damages of even neuronal hyperactivity are able to recruit and activate microglia, even in the absence of cell death and leakage of the intracellular ATP (Eyo et al., 2014). Third, microglia can rapidly extend processes within minutes after injury by sensing ATP, while the activation of cellular apoptotic pathway usually takes more than 30 minutes to hours (Rehm et al., 2003). Last but not least, the level of microglia migration can be modulated by the injury strength, while apoptotic relevant ATP release happens in an all-or none type that lacks precise regulation. Altogether, there should exist an alternative source and mechanism for controlling ATP release, that can both rapidly respond after injury, as well as maintain effective ATP concentration in time and space for guiding microglia.

In our previous effort, we advanced the technique for sensitive monitoring ATP dynamics by engineering novel genetically-encoded fluorescent sensors GRAB_ATP1.0_ (ATP1.0), and demonstrated its utility in tracking extracellular ATP in vivo (Wu et al., 2022). Here, equipped with such advanced tool, we further combined in vivo two-photon imaging and laser injury model, to precisely tracked the ATP events at sub-second and cellular resolution, and across the entire injury process that matches the microglial response. Together with mechanistic studies, we uncovered how ATP is able to encode injury information and regulates the complex brain injury response.

## Results

### Injury-evoked ATP exhibits characteristic spatiotemporal patterns

To quantitatively assess the brain internal response to injury, we established a focal laser ablation (FLA) model (Davalos et al., 2005; Galbraith and Terasaki, 2003) to precisely introduce injury *in vivo*, and recorded microglial mobility as a primary readout of the injury response. In this model, brief (2 s) and local (50 μm in diameter) injury mobilized microglia from a broad area (even hundreds of micrometers away from the injury) to carry out directional process extension and migration that lasted for hours (Figure S1A; simply as ‘migration’ below). The spatiotemporal difference between the external injury and internal microglial migration suggested that the microglia themselves may not be the element that accurately senses injury; rather, they may indirectly perceive injury that was presented by other cells or molecules. In principle, this “injury-presenting” signal should bridge the injury and the microglial response by both encoding the injury properties, and exhibiting similar patterns with microglial migration in time and space to provide guidance cues.

To check whether extracellular ATP, a critical molecule in regulating microglial migration (Davalos et al., 2005; Shizuyo et al., 2001), could act as the intermediate signal, we virally expressed the recently developed ATP1.0 sensor in either astrocytes or neurons to monitor ATP dynamics extracellularly during FLA (Figures 1A, S1B and S1C). After FLA, two distinct types of ATP dynamics were observed, including a strong burst-like increase in fluorescence that surrounded the injury center (ATP burst), and multiple flashing pattern of events across the entire field (Figures 1B–D, S1C and Video S1). The flashing type of ATP release was observed for the first time *in vivo* with the high sensitivity and spatiotemporal resolution of ATP sensor. We referred to the individual flashing event as an <u>Inflare</u>, which is short for <u>in</u>jury-induced <u>fla</u>shing ATP <u>re</u>lease. Both signal types were identified when ATP1.0 was expressed by either astrocytes or neurons, with astrocytic expression showing more robust signals (Figures S1C– S1I), and were selected for the following experiments.

**Figure 1.**
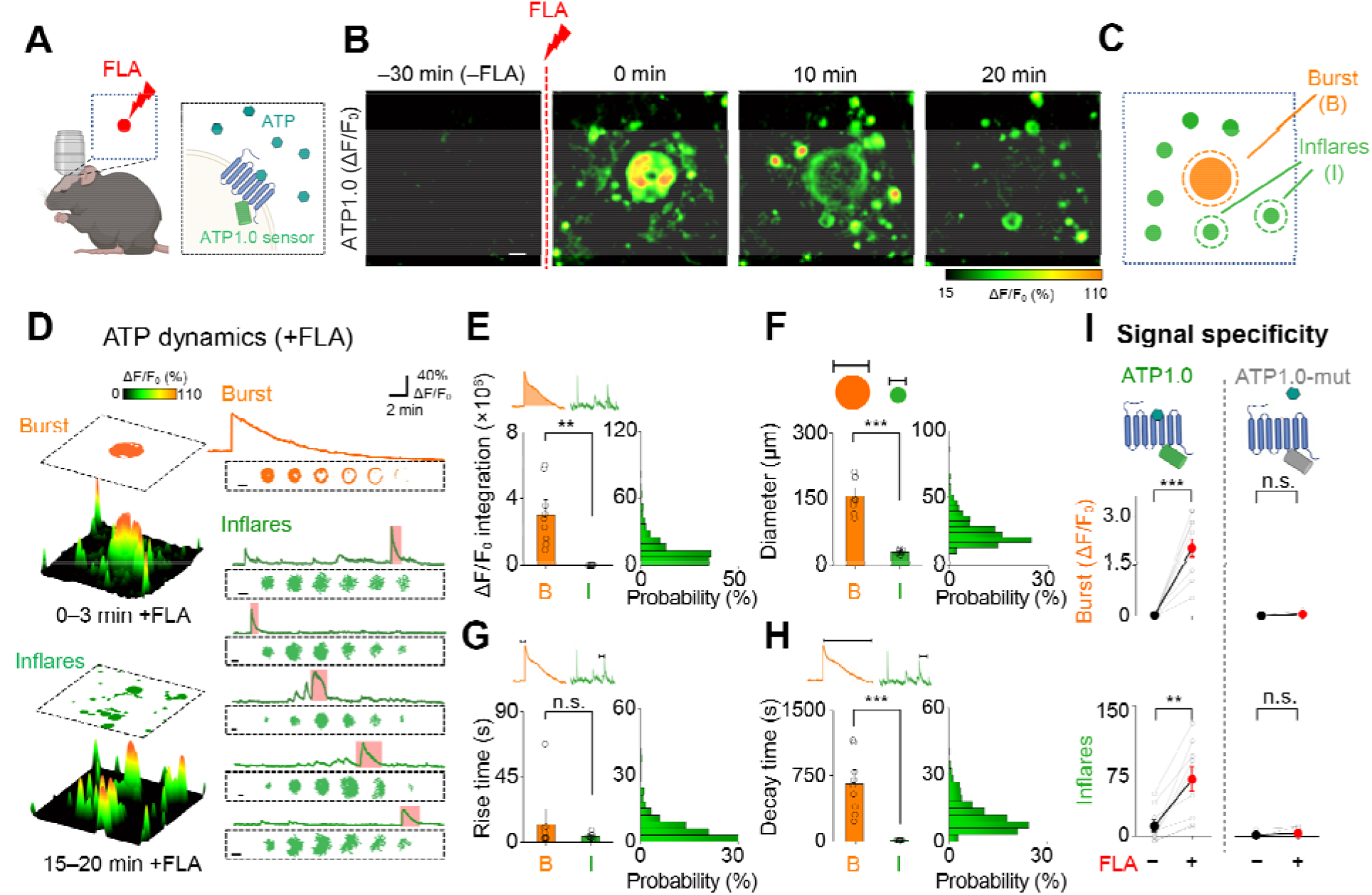
Direct observation of ATP dynamics *in vivo* during brain injury. **(A)** Schematic of FLA injury during two-photon imaging in mice (left), and the fluorescent sensor ATP1.0, which is an indicator of extracellular ATP dynamics (right). **(B)** Fluorescence response (shown as ΔF/F_0_ pseudo-color) of ATP1.0 before (−FLA) and after FLA at the indicated time points. **(C)** Cartoon of the two types of ATP events. **(D)** Representative regions of interest (ROIs) showing an ATP burst and Inflares identified from the fluorescence response after FLA (+FLA). Traces of an ATP burst and Inflares are shown, with the progression of Inflares over space (during the time indicated by shaded areas) shown below each trace. (**E–H**) Comparison of the fluorescence response integration (**E**), diameter (**F**), rise time (**G**) and decay time (**H**) between ATP bursts and Inflares. Group data from each mouse are shown on the left, with the distribution of individual Inflares summarized on the right (*n* = 9 mice, with *n* = 9 ATP bursts and *n* = 643 Inflares quantified). (**I**) Group analysis of the fluorescence response of ATP bursts (upper) and the number of Inflares (lower, over 20 min) in mice expressing ATP1.0 (green, left) or ATP1.0-mut (gray, right) after FLA (*n* = 9 and 8 mice, respectively). Scale bars, 50 μm. Data are shown as the mean ± s.e.m. \*\**p* < 0.01, \*\*\**p* < 0.001; n.s., not significant. See Table S1 for statistics.

Quantification indicated that ATP bursts and Inflares could be clearly distinguished based on signal properties—specifically, the ATP burst was characterized by a higher amplitude in the fluorescence response (ΔF/F_0_), a larger diameter and longer duration (slower on/off kinetics) as compared with individual Inflares (Figures 1E–1H). Also, the ATP burst was immediately and uniquely presented after each injury, whereas Inflares were generated with a frequency of 2–3 events/min (over a 20 min recording after FLA; Figure 1I). We verified that all responses represented specific ATP release rather than autofluorescence, as expressing an ATP1.0-mutant sensor that lacks the ATP-binding ability showed no fluorescence changes after FLA (Figures 1I and S2A). Also, these spatially-selective Inflares did not result from heterogeneous sensor expression, as no correlation was found between basal fluorescence and signal changes (Figures 2B–2D). Among Inflares, the properties of individual events were largely homogeneous, even though they were generated at different times and locations (Figure S3). Together, using new fluorescent sensors *in vivo*, we characterized two distinct types of ATP dynamics with high spatiotemporal selectivity after brain injury.

**Figure 2.**
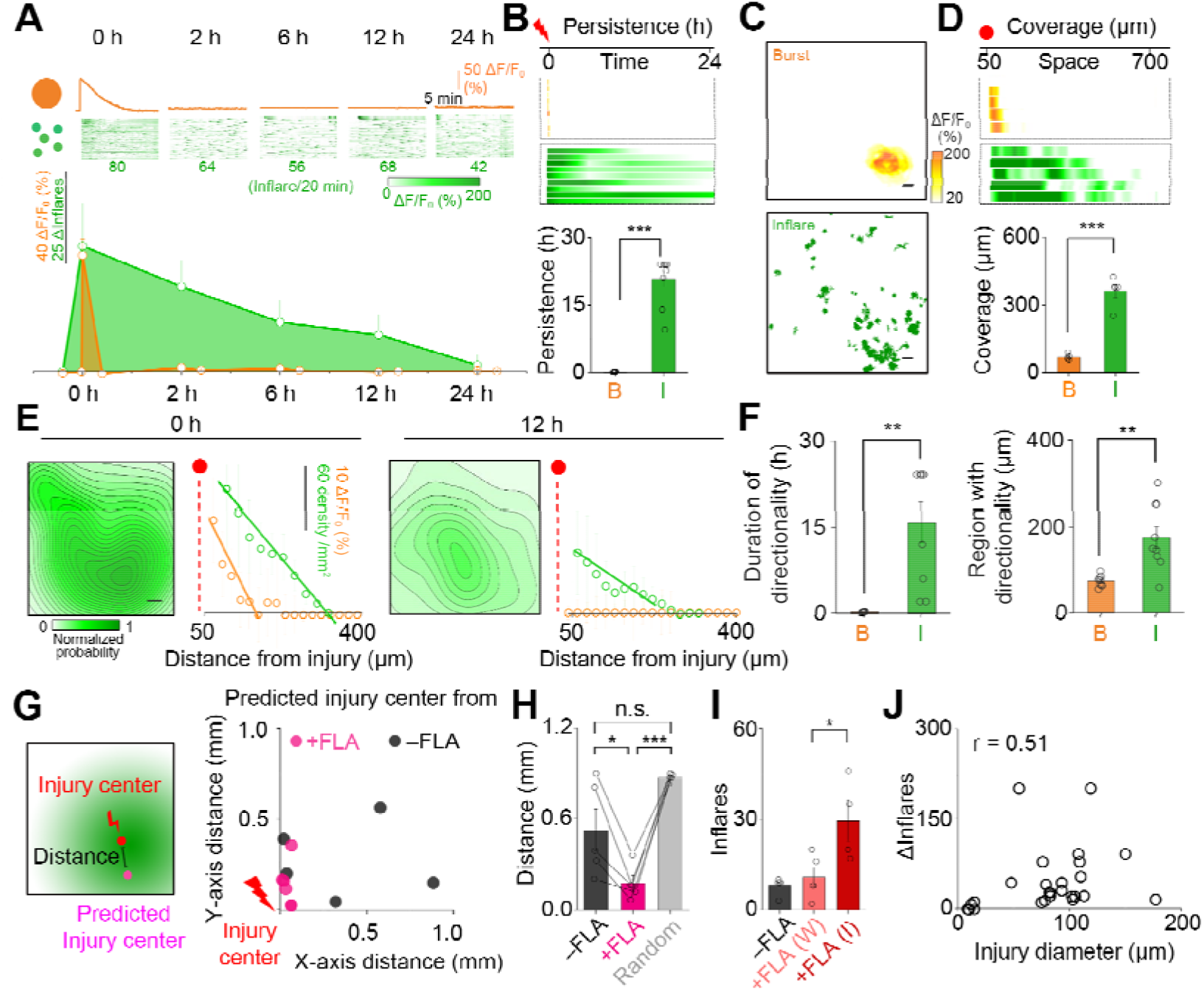
Persistent and widespread Inflares encode injury information. **(A)** Temporal changes in ATP signals over a 24 h period after injury. Representative traces of ATP bursts and Inflares at the indicated times are shown (upper, each line represents individual Inflares, with the number of Inflares labeled below), with the averaged signal (ΔF/F_0_ for the burst and number for the Inflares) quantified (lower; *n* = 9 mice). **(B)** Persistence of two types of ATP signals (*n* = 9 mice), with each line representing one mouse. **(C)** Representative image showing the spatial distribution of ATP bursts and Inflares after injury (upper, ATP bust; lower, Inflares). **(D)** Coverage of two types of ATP signals (*n* = 5 mice), with each line representing one mouse. **(E)** Contour map of the inflare frequency gradient at indicated times after injury, and quantification of the gradient distribution generated by Inflares or ATP bursts (*n* = 5 mice). **(F)** Duration of directionality (left) and region with directionality (right) of the ATP bursts and Inflares after FLA injury (*n* = 9 mice). The region with directionality was quantified as the cross point of the fitted gradient curve (shown in E) with the distance to injury. **(G)** Left, schematic of the predicted injury center based on the Inflare frequency gradient, and the distance between this predicted center to the FLA injury center. Right, the location of the predicted injury center based on the Inflare distribution before (–FLA) and after (+FLA) injury, with the position of the real injury center set as the origin (*n* = 5 mice). **(H)** Distance between the Inflare-predicted injury center and the FLA injury center. Inflares recorded before FLA or artificial generated events with same frequency but randomly distributed in space (Random) were used as controls (*n* = 5 mice). **(I)** Inflare number after weak (W) or intensive (I) FLA injury (*n* = 5, 5 and 4 mice for no FLA (**−**FLA), weak and intensive FLA, respectively). The weak and intensive FLAs were conducted by setting the laser power at 60% for 0.02 s and 100% for 4 s, respectively. **(J)** Correlation between injury diameter and changes in Inflare number (ΔInflares) after injury (*n* = 28 mice in total). The injury diameter was measured based on the damaged region in images after FLA. Scale bars, 50 μm. Data are shown as the mean ± s.e.m. \**p* < 0.05, \*\**p* < 0.01, \*\*\**p* < 0.001; n.s., not significant. See Table S1 for statistics.

### Inflares encode injury information and persistently guide microglial migration

With specific ATP dynamics recorded, we next determined whether they could encode and present injury information to microglia. As microglia show persistent response across a broad region after injury, we first mapped the changes of ATP events across time and space to see whether they shared similar properties with microglia. Interestingly, although the ATP burst and Inflares were both injury-responsive events, their correlation with injury differed greatly. For quantitative comparison, we defined the time of effective signal presentation (higher than baseline without FLA) as signal persistence, and the maximum distance of effective signal presentation from injury as signal coverage. The persistence of total Inflares (∼20.8 h) was over 100-fold longer than that of the ATP burst (∼0.17 h) (Figures 2A, 2B and S4), and the coverage of Inflares (∼360 μm) was fivefold greater than that of the ATP bursts (∼70 μm) (Figures 2B and 2D). Within the covered area, the Inflare frequency stepped down with increasing distance from the injury site, thus generating an “ATP event gradient” across a broad region (>300 μm), which could be maintained for hours after injury (Figures 2E and 2F). In contrast, the ATP burst by its concentration exhibited a limited spatial gradient, which ended with the disappearance of the signal after 20 min (Figures 2A, 2E and 2F). Therefore, the spatiotemporal properties of Inflares, rather than of the ATP bursts, were closely matched with the characteristics of microglial migration.

As the initial response to injury, ATP signals might encode injury information that is critical for guiding following response. To explore whether injury information can be extracted from properties of Inflares, we first predicted the injury center from the frequency gradient of Inflares, and compared the predicted injury center with the FLA site applied. The Inflare predicted injury center matched closely with the FLA sites (Figure 2G). In sharp contrast, the prediction from neither Inflares that were recorded before FLA nor randomly distributed artificial events with a similar frequency matched with the injury sites (Figure 2H), supporting that Inflares could encode positional information of injury. In addition, Inflares also encoded injury strength, as their frequency positively correlated with FLA intensity and damaged size (Figures 2I and 2J). Interestingly, other properties within individual Inflares, including the size and duration did not correlate with injury strength, suggesting the encoding of injury by Inflares existed at group rather than individual level (Figure S5). Altogether, these results implied that Inflares were able to encode injury information in multiple dimensions.

To directly test the contribution of Inflares, especially persistent ones, on presenting injury to guide microglia migration, we performed a temporally specific perturbation of ATP by injecting the ATP degradation enzyme apyrase (Dubyak and el-Moatassim, 1993) into the cortex, either prior to the injury (pre-injury) to block all ATP signals, or 30 min after injury (post-injury) to selectively block persistent Inflares afterwards (Figures 3A–3C). ATP1.0 imaging verified that pre-injury injection of apyrase substantially decreased ATP bursts and Inflares, whereas the post-injury injection of apyrase left the ATP burst unaltered but selectively inhibit Inflares and reduced their persistence. Injection of artificial cerebrospinal fluid (ACSF) as controls (either pre- or post-injury) did not affect the ATP signals (Figures 3D–3F). Next, the same protocol was repeated in microglia labeling Cx3cr1-GFP mice to monitor changes of microglial migration.

**Figure 3.**
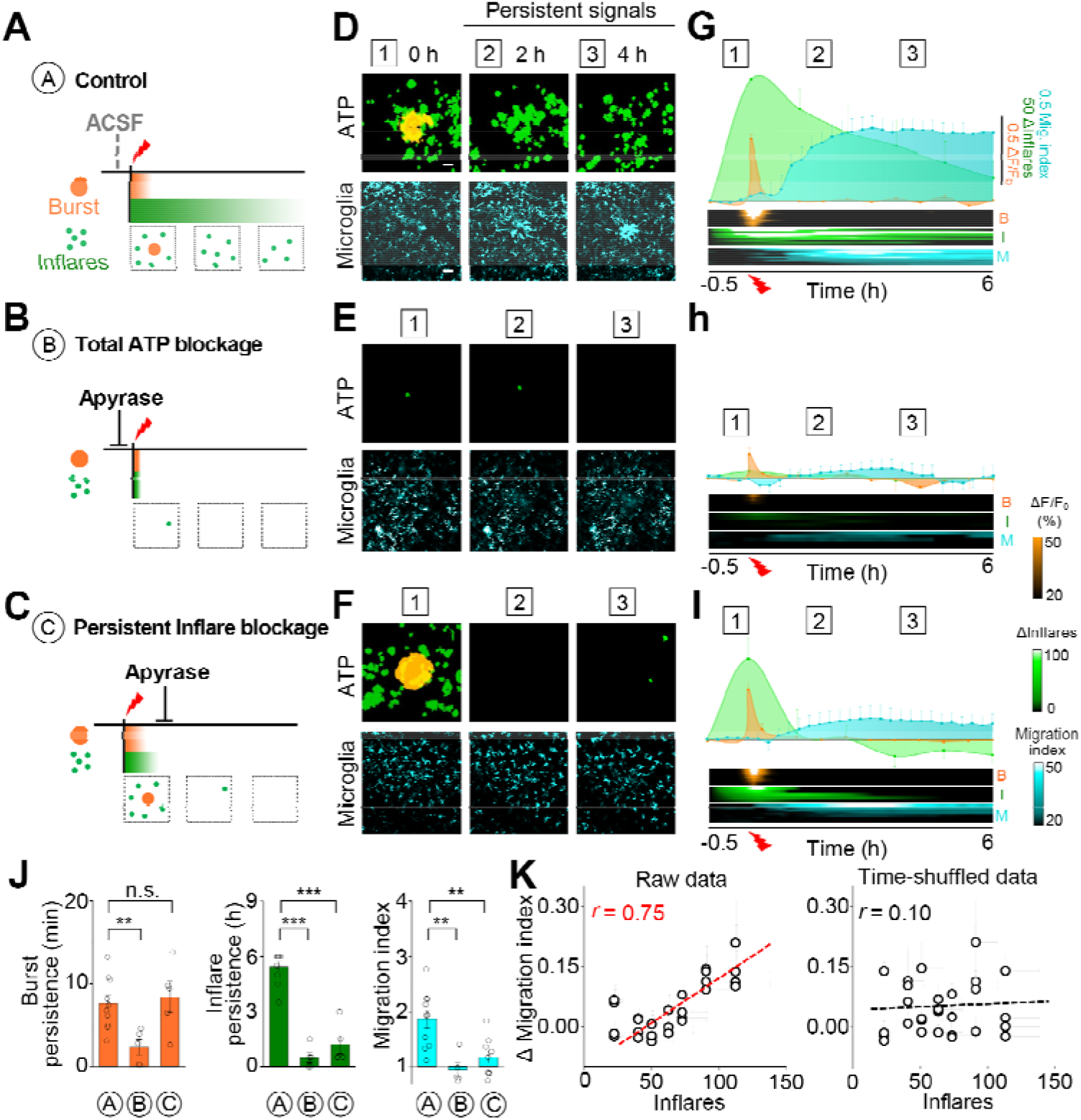
Inflares rather than the ATP burst drive microglial migration. (**A–C**) Illustration of the temporally specific perturbation of ATP events by local apyrase injection. The same volume of ACSF was injected as a control. (**D–F**) Representative images showing the ATP dynamics (upper) and microglial migration (lower) at the indicated time points after injury. ROIs of the ATP bursts and Inflares were identified and merged. (**G–I**) Changes in ATP signals (ΔF/F_0_ for the ATP burst and the number of Inflares) and microglial migration after injury. Raw data from individual mice are shown at the bottom, with each row representing one mouse (B: ATP burst; I: Inflares; M: microglia migration). **(J)** Group analysis of the ATP signal persistence (*n* = 10, 5, 5 mice in group A, B, C, respectively) and index of microglial migration (*n* = 10, 5, 9 mice in group A, B, C, respectively). **(K)** Correlation between the microglial migration index and the number of Inflares after injury. Raw data (left) and time shuffled data (right) are shown with the Pearson correlation coefficient indicated. Scale bars, 50 μm. Data are shown as the mean ± s.e.m. \*\**p* < 0.01; \*\*\**p* < 0.001; n.s., not significant. See Table S1 for statistics.

Blocking all ATP signals caused no observable microglial migration after FLA, in contrast to robust and sustained migration in the ACSF-injected mice (Figures 3G and 3H). Crucially, selectively reducing the persistence of Inflares, without affecting the ATP burst, also led to inhibition of migration (Figures 3I and 3J). These changes in Inflares and microglia were not caused by other factors, including differences in injury, variability in sensor expression, or levels of microglial activation across experiments (Figures S6A–S6J). In addition, heat-inactivated apyrase, which lost the enzyme activity to degrade ATP (Walter et al., 1951), was not able to inhibit microglial migration, confirming the dependence of its effect on ATP levels (Madry et al., 2018) (Figure S6K). Furthermore, a strong positive correlation was found between Inflare frequency and changes in microglial migration at each recording period, which was not seen with time-shuffled pairing or between ATP burst and microglial migration (Figures 3K and S6L). We therefore concluded that persistent Inflares rather than ATP bursts were key signals that drove microglial migration toward the site of injury.

### Astrocytic Ca^2+^ activity generates Inflares through pannexin 1 channels

The persistent generation of Inflares from multiple locations suggested that they might have a unique regulatory mechanism at the cellular and molecular level. Although multiple cell types have been reported to actively release ATP (Guthrie et al., 1999; Imura et al., 2013; Sperlágh and Vizi, 1996), most of these studies were carried out in cultured cells which may not represent complex physiological regulation *in vivo*. To uncover the cellular source of Inflares, we manipulated individual cell types by either ablation or inhibition, and checked their effect on FLA-induced Inflares *in vivo*. Selective ablation of local astrocytes was achieved by either injection of the gliotoxin L-α-aminoadipate (L-α-AA) (Khurgel et al., 1996) or astrocyte-specific expression of herpes simplex virus thymidine kinase (HSV-TK) followed by ganciclovir (GCV) delivery (Delaney et al., 1996). With either strategy, no FLA-evoked increases in Inflares were observed (Figures 4A–4C, and S7A–S7D; in these experiments, ATP1.0 was expressed in neurons). In contrast, ablation of neurons by selective caspase-3 expression (AAV-hSyn-taCasp3) (Yang et al., 2013) or silencing of neural activity by expressing an engineered human muscarinic M_4_ receptor (hM_4_Di) with clozapine-N-oxide (CNO) delivery did not affect FLA-evoked Inflares (Figures 4D, 4E, and S7E–S7H). FLA was also able to evoke a robust increase in Inflares when microglia were depleted by the CSF1R inhibitor, BLZ945 (Hagemeyer et al., 2017) (Figures 4F, 4G, S7I and S7J). These results showed that astrocytes were the major cellular source of Inflares.

**Figure 4.**
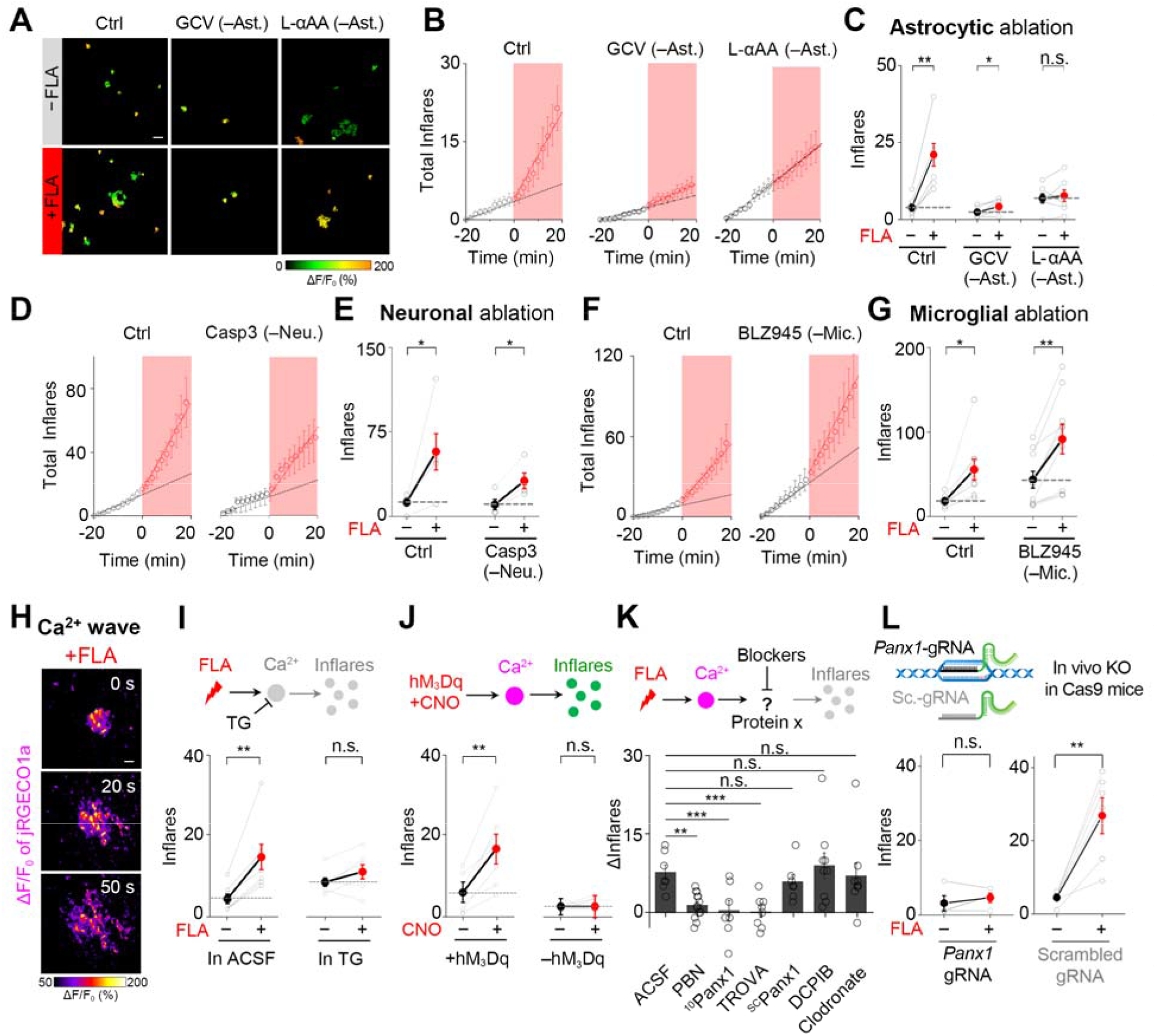
Astrocytes selectively generate Inflares that depend on intracellular Ca^2+^ and pannexin channels. (**A**) Representative images of Inflares (shown as ROIs) before and after FLA injury when astrocytes were selectively ablated. (**B, C**) Changes in the total number of Inflares (B) and group analysis of Inflare number (C) when astrocytes were selectively ablated. Mice expressing HSV-TK without the application of GCV were used as control (Ctrl) (*n* = 6, 6 and 7 mice for the Ctrl, GCV and L-αAA groups, respectively). Time of FLA injury was set as 0. Dashed lines indicate the predicted number of Inflares based on the baseline frequency. **(D)** Changes in the total number of Inflares when neurons were ablated (*n* = 5 and 4 mice for Ctrl and Casp3, respectively). Mice with neuronal expression of mCherry were used as control (Ctrl). **(E)** Group analysis of Inflare number when neurons were ablated (*n* = 5 and 4 mice for Ctrl and Casp3, respectively). (**F, G**) Similar to (D, E), except microglia were selectively ablated by oral administration of BLZ945 (*n* = 8 and 9 mice for Ctrl and BLZ945, respectively). BLZ945 was added in the food pellet (see Methods for detail), and mice fed with vehicle (saline)-added food were used as control (Ctrl). **(H)** Representative Ca^2+^ waves shown by the fluorescence signal (ΔF/F_0_) of astrocytic expressing jRGECO1a at the indicated time after FLA. **(I)** Inflare number in ACSF- or thapsigargin (TG)-treated slices after FLA (*n* = 7 slices from 3 mice in each group). **(J)** Number of Inflares in ACSF- and CNO-treated slices with or without hM_3_Dq expression (*n* = 7 slices from 3 mice (7/3) for +hM_3_Dq, *n* = 8/2 for −hM_3_Dq). **(K)** Changes of Inflare number in brain slices in the presence of the indicated blockers (*n* = 7 slices from 3 mice for ACSF, *n* = 13/4 for PBN, *n* = 7/3 for ^10^panx1, *n* = 8/3 for TROVA, *n* = 7/3 for ^sc^panx1, *n* = 9/3 for DCPIB, *n* = 9/3 for clodronate). **(L)** Inflare number in response to FLA in astrocytic *Panx1* gene knockout mice *in vivo* (*n* = 4 mice). *Panx1* gene knockout was achieved by injecting AAVs expressing gRNAs (under the astrocyte-specific promoter GfaABC1D) for *Panx1* in LSL-Cas9 transgenic mice. A scrambled gRNA (sc.-gRNA) was used as the control. Scale bars, 50 μm. Data are shown as the mean ± s.e.m. \**p* < 0.05; \*\**p* < 0.01; \*\*\**p* < 0.001; n.s., not significant. See Table S1 for statistics.

We next explored whether injury-induced changes in astrocytic activities caused the generation of Inflares. Brain slices from mice expressing the red Ca^2+^ sensor jRGECO1a (Dana et al., 2016) and ATP1.0 were used to record astrocytic Ca^2+^ activity and Inflares simultaneously. Immediately after FLA, a continuous Ca^2+^ wave within astrocytes was observed, which originated from the injury center and further propagated across the entire imaging field (Figures 4H and S8A). Such Ca^2+^ waves are not likely to be the downstream of the ATP burst, as the application of broad ATP receptor antagonists (blocked the effect of ATP) did not affect the generation of the astrocytic Ca^2+^ wave (Figure S8B). We further explored the causal relationship between this Ca^2+^ wave and Inflares. Depleting intracellular Ca^2+^ with the SERCA inhibitor thapsigargin (TG) resulted in no Inflare increase after FLA (Sehgal et al., 2017) (Figures 4I and S8A), and blocking the gap junction with carbenoxolone (Rozental et al., 2001) (CBX) reduced Ca^2+^ wave propagation and decreased the coverage of Inflares (Figures S8C and S8D), indicating that both the generation and propagation of Ca^2+^ waves were necessary for Inflares. To test the sufficiency of Ca^2+^, we elevated astrocytic Ca^2+^ by a chemogenetic approach in the absence of injury, in which the application of CNO could lead to an increase in intracellular Ca^2+^ by activating the hM_3_Dq receptor (Agulhon et al., 2013). CNO application in hM_3_Dq-expressing astrocytes significantly increased Inflares, whereas CNO alone did not (Figures 4J and S8E). Astrocytes with chemogenetically elevated Ca^2+^ shared a similar activation status as did those after injury, as further introducing FLA in the presence of CNO failed to evoke an additional increase in Inflares (Figure S8F).

To identify the molecular mechanism responsible for Inflares, we identified candidate proteins previously shown to participate in active ATP release (Hyzinski-García et al., 2014; Iglesias et al., 2009; Oya et al., 2013) and that are highly expressed in astrocytes (Chai et al., 2017). Selective pharmacological blockers for these candidates were individually applied in brain slices to test their effects on FLA-evoked Inflares. Multiple blockers of the pannexin 1 hemichannel, including probenecid (PBN) (Silverman et al., 2008), trovafloxacin (TROVA) (Poon et al., 2014) and peptide inhibitor ^10^panx1 (Brough et al., 2009) were all effective in blocking FLA-induced Inflares, whereas the scrambled control peptide (^sc^panx1) did not (Figures 4K and S8G).

Inhibitors for other candidates such as the volume-regulated anion channel (VRAC) (Qiu et al., 2014) and the vesicular nucleotide transporter (vNUT) (Kato et al., 2017) failed to block Inflares (Figure S8G). Importantly, knockout of the gene encoding pannexin 1 specifically in astrocytes, in either transgenic mice (*Panx1*^flox/flox^ injected with AAV-GFaABC1d-cre) or by an AAV-mediated Cas9-gRNA strategy also resulted in no increase in Inflares after FLA (Figures 4L and S8H), verifying the pharmacological screening results. Consistently, CNO-induced Inflares in hM_3_Dq-expressing astrocytes were also blocked by the pannexin inhibitor PBN (Figure S8I). In conclusion, injury evoked Inflares were generated in an astrocytic Ca^2+^ wave–dependent manner through the opening of the pannexin 1 hemichannel.

### Inflares drive broad changes in microglial activities and are potential therapeutic targets for stroke

In addition to directional migration, microglia are able to dynamically tune their broad range of activities in response to ATP (D Ferrari et al., 1996; Davide et al., 1999; McLARNON. et al., 1999), and consequently affect injury management and outcome. By selectively targeting pannexin 1, we examined the downstream effect of Inflares on microglial function and injury status. In addition to guiding migration, Inflares greatly promoted the morphological transformation of microglia around the injury site, which was inhibited by the pannexin blocker PBN (Figures 5A, 5B and S9A). Microglia surrounding the injury also showed increased proliferation only when Inflares were present (Figure 5C). Functionally, Inflares were necessary to promote microglial phagocytosis of apoptotic cells within the region of the injury (Figure 5D). We also checked the expression of key mediators for immune response and injury repairing 24 h after extensive FLA injury. The presence of Inflares boosted pro-inflammatory gene expression including *Il1b* and *Cxcl10* and at the same time reduced both the anti-inflammatory genes (such as *Il10* and *Tgfb1*) and several neurotrophic factors. Treatment with PBN or astrocytic knockout of *Panx1* returned the injury-induced transcriptional changes to resting levels (Figures 5E, F and S9C–E). These results indicated that injury broadly affected microglial functions, effects that were critically dependent on Inflares. Consequently, the dysfunctional microglia induced by injury further increased the local cellular apoptosis in an Inflare-dependent manner, as shown by TUNEL-positive cells and the expression of activated caspase-3 after injury (Figure 5G).

**Figure 5.**
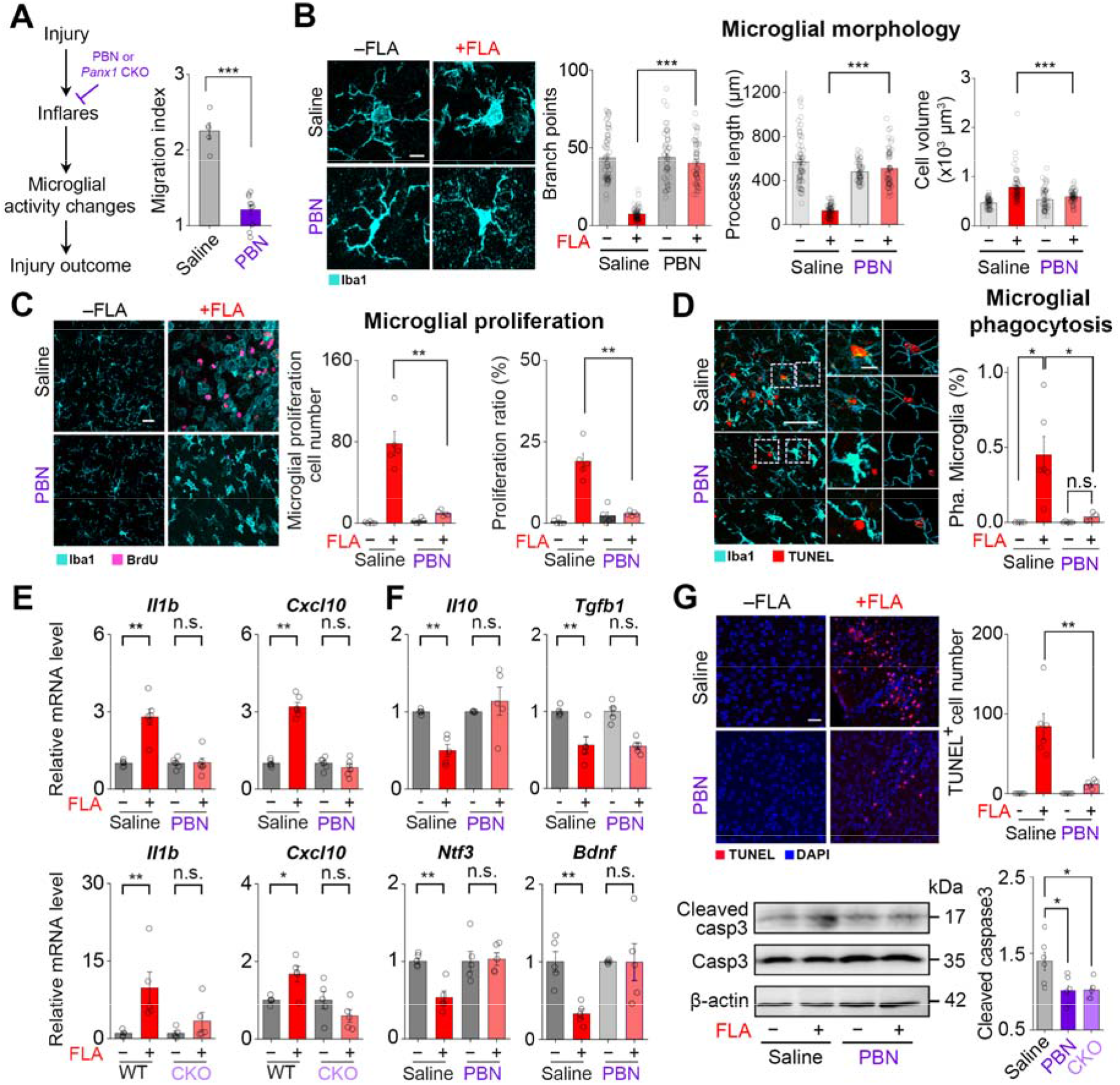
Inflares alter multiple aspects of microglial activity and exacerbate local damage. **(A)** Changes in microglia migration *in vivo* in saline or PBN-treated mice (8 h after FLA; *n* = 5 in saline group, *n* = 10 in PBN group). **(B)** Morphological changes in microglia (stained with Iba1) in saline- and PBN-treated mice with or without FLA. For branch point data, saline: *n* = 63/6 cells/mice for −FLA, *n* = 63/6 for +FLA; PBN: *n* = 52/5 for –FLA, *n* = 55/6 for +FLA. For process length, saline: *n* = 63/6 cells/mice for −FLA, *n* = 63/6 for +FLA; PBN: *n* = 52/5 for −FLA, *n* = 55/6 for +FLA; for cell volume, *n* = 61/6 cells/mice in each group. **(C)** Representative images and quantification of injury-induced microglial proliferation after 48 h in saline- or PBN-treated mice stained with BrdU and Iba1 (*n* = 5 mice in each group). The proliferating microglia were counted based on the colocalization between BrdU^+^ and Iba1^+^ signals, and the proliferation ratio was defined as proliferating microglia over total microglia. **(D)** Representative images (left, +FLA) and quantification of microglial phagocytosis before (– FLA) and 8 h after (+FLA) injury in saline- and PBN-treated mice. Phagocytosis was identified by colocalization of TUNEL-positive (TUNEL^+^) apoptotic cells together with Iba1^+^, and the fraction of phagocytosing microglia over total microglia was quantified (saline: *n =* 4 mice for – FLA, *n* = 6 for +FLA; PBN: *n* = 5 for –FLA, *n* = 4 for +FLA). Regions in the dashed boxes are zoomed in on the right. **(F)** Cytokine gene expression in saline- and PBN-treated mice (upper) and in astrocytic *Panx1* knockout mice (CKO; lower) at 24 h after FLA. Data for additional genes are summarized in Figure S9 (*n* = 6 mice for saline and PBN, *n* = 5 for wild type (WT) and CKO). **(F)** Anti-inflammatory (upper) and neurotrophic (lower) gene expression level in saline- and PBN-treated mice at 24 h after FLA (*n* = 5 mice in each group). **(G)** Cell apoptosis at the injury site 8 h after FLA, quantified by TUNEL^+^ cell numbers (upper; *n* = 6 mice) and the activation of caspase-3 (lower; cleaved caspase-3 relative to total caspase-3; *n* = 6 mice for saline, *n* = 6 for PBN, *n* = 5 for CKO). Scale bars, 50 μm, except 10 μm in (B) and in zoomed images in (D). Data are shown as the mean ± s.e.m. \**p* < 0.05; \*\**p* < 0.01; \*\*\**p* < 0.001; n.s., not significant. See Table S1 for statistics.

Finally, to assess the contribution of Inflares at the animal level, we performed middle cerebral artery occlusion (MCAO) in mice to mimic ischemic stroke (Howells et al., 2010) (Figure 6A). Strikingly, 12 h after MCAO reperfusion, intensive Inflares were observed in the cortex, with the frequency reaching ∼20 events/min (10 times higher than the rate associated with FLA; Figure 6B and Video S2). In contrast, Inflares were rarely recorded in sham surgery mice (Sham) or in mice expressing the ATP1.0-mutant sensor and subjected to MCAO (Figure 6B). Individual Inflares had properties similar to those after FLA injury, suggesting a common cellular and molecular mechanism (Figure 6C). Indeed, blocking the pannexin channel by PBN led to no increase in Inflares after MCAO (Figures 6B and 6D). Functionally, blocking Inflares prevented pathological changes in local transcription after MCAO (Figures 6E and S10), and area of brain infarction was significantly reduced, especially in the penumbra cortical region which faced secondary injury after stroke (Figure 6F). As a consequence of reduced brain damage, mice with Inflares blockage recovered more effectively after stroke with minor behavioral abnormalities, including lower neurological deficit scores, stronger locomotion and balancing ability as compared with MCAO mice without treatment (Figures 6G–I). Thus, the key molecule pannexin 1, which is responsible for Inflares, represents a promising drug target for reducing stroke-related secondary injury and promoting disease outcome.

**Figure 6.**
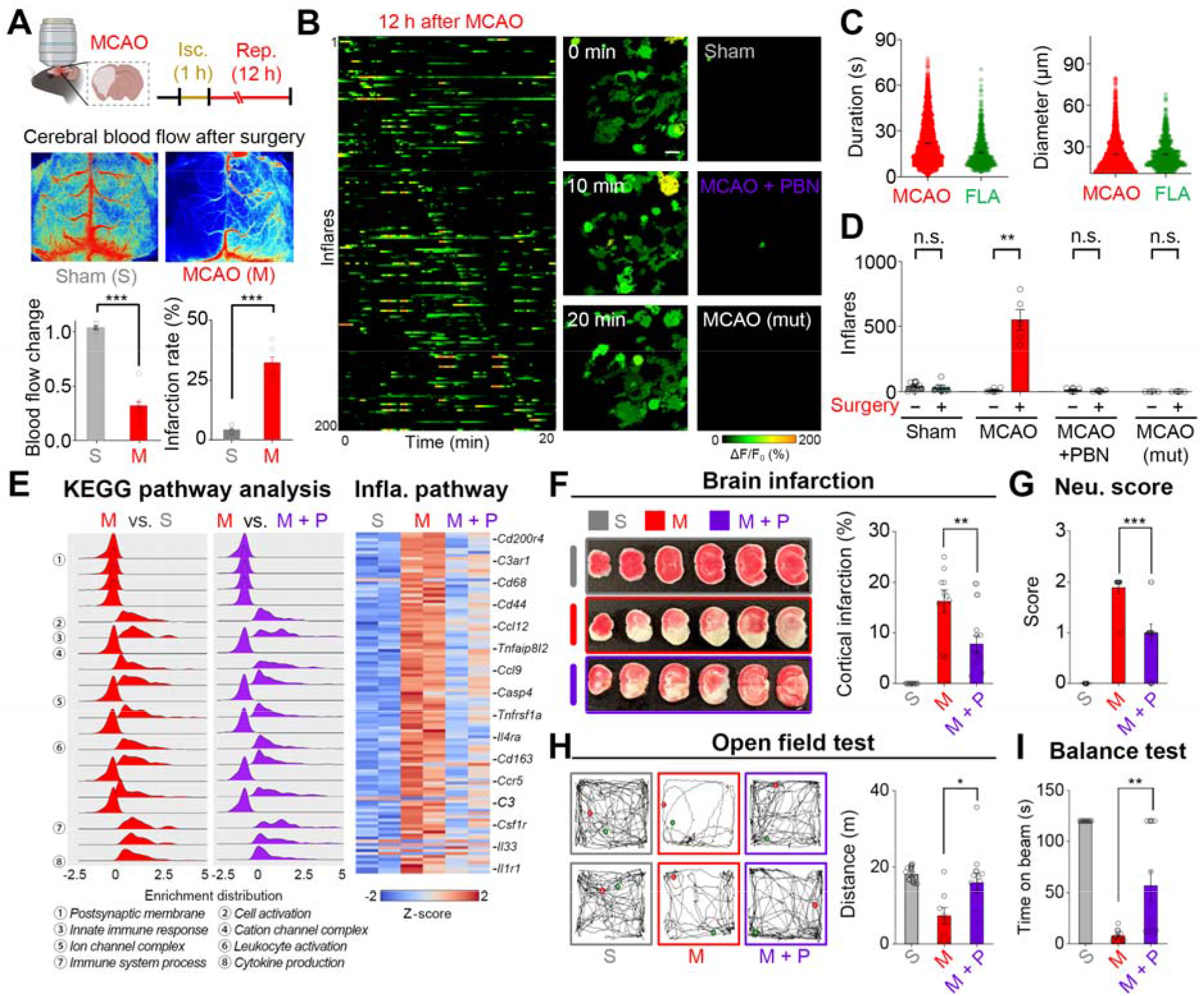
Blocking excessive Inflares by targeting pannexin 1 benefits stroke outcome. **(A)** The MCAO mouse model, with 1 h of ischemia (Isc.) followed by 12 h of reperfusion (Rep.), showed hemisphere-specific reduction of blood flow (*n* = 5 and 9 mice for sham and MCAO, respectively), and an increase in brain infarction quantified from triphenyltetrazolium chloride (TTC) staining results (*n* = 6 mice in each group). **(B)** Left, representative traces of Inflares at 12 h after reperfusion in MCAO mice. Middle and right, ROIs with Inflares at the indicated time points in MCAO, sham surgery and MCAO with PBN treatment (MCAO + PBN) mice and in MCAO mice expressing ATP1.0-mut (MCAO (mut)). **(C)** The duration and diameters of individual Inflares recorded after MCAO (red) or FLA (green) (*n* = 2913/6 ROIs/mice for MCAO, *n* = 643/9 for FLA). **(D)** Group analysis of Inflare numbers (*n* = 6 mice for sham, *n* = 5 mice for MCAO, *n* = 4 mice for MCAO + PBN, *n* = 4 mice for MCAO (mut)). **(E)** KEGG pathway analysis of the differentially expressed genes when comparing MCAO with sham (left) or MCAO with MCAO + PBN samples (middle). Expression profiling of the inflammatory pathway plotted as a heat map of Z-scores (right). **(F)** Representative TTC staining images (left) and group analysis of the cortical infarction rate 3 days after surgery in sham, MCAO and MCAO + PBN mice (*n* = 10, 9 and 12 mice, respectively). **(G)** The neurological deficit scores (Neu. score) for sham, MCAO and MCAO + PBN mice evaluated 3 days after surgery (*n* = 10, 9 and 12 mice, respectively). **(H)** The locomotion ability of sham, MCAO and MCAO + PBN mice in an open-field test 3 days after surgery (*n* = 10, 8 and 12 mice, respectively), with the trajectory of two representative mice in each group shown on the left. **(I)** Amount of time that mice stayed on the balance beam in sham, MCAO and MCAO + PBN mice (*n* = 10, 9 and 12, respectively). Scale bars, 50 μm. Data are shown as the mean ± s.e.m. \**p* < 0.05; \*\**p* < 0.01; \*\*\**p* < 0.001; n.s., not significant. See Table S1 for statistics.

## Discussion

In contrast to the well-established role of purinergic transmitters in recruiting microglia, and the precise characterization of microglia motility after injury, no evidence so far has directly reported the dynamics of ATP signal as well as its regulation in vivo. In this study, we directly observed a characteristic pattern of ATP dynamics, Inflares, that serves as injury-responsive signals. In contrast to the previous notion that ATP only released from damaged apoptotic cells passively after injury, we found that Inflares were highly regulated events that are dependent on intracellular Ca^2+^ and pannexin hemichannels in astrocytes. Our finding illustrated the functional cooperation between cells for the complex injury response, in which astrocytes were specialized to sense injury, and presented injury by Inflares to activate microglia for injury management.

Such functional cooperation increased the sensitivity and efficiency of the injury response, as microglia can effectively mediate the injury response but are limited in number with isolated territories (Kettenmann et al., 2011), their direct detection of injury might lead to missed signals, whereas astrocytes, which form a tight functional network, are more robust in detecting injury and amplifying signals.

The precise identification and presentation of injury, including its intensity and position, are a prerequisite for proper injury response. Here we found that Inflares could encode injury information by their properties at group level, thus faithfully guide microglia to the injury site. Instead of using an absolute concentration gradient, Inflares encoded direction based on their frequency distribution, which increased the robustness of information delivery under the effective degradation of ATP *in vivo* (Kege et al., 1997; Zimmermann, 2000). Similarly, the “Inflare mode” of ATP release, compared with a “one-shot” releasing mode, represents an elegant design for maintaining a sustained presence of ATP even in the context of effective degradation, as well as expanding the effective coverage of the ATP signal in space. Consequently, the temporal and spatial amplification achieved by Inflares provides a time window during which slow cellular processes such as migration can occur, and allows the recruitment of responsive cells from a broader area to ensure the proper management of injury.

The activity changes of microglia have been reported to exert multifaceted effects that were highly dynamic and context dependent, which contributed to their complex roles in the brain (Colonna and Butovsky, 2017; Nayak et al., 2014). Therefore, although changes in microglia have been associated with multiple diseases, it might be ineffective to isolate or manipulate only their individual activities or functions for therapeutic purposes. By blocking Inflares, which acted as the upstream signal for changes in microglial behaviors, we successfully reversed the dysfunction of microglia after injury across multiple aspects, which suggests their potential as a target for preventing secondary damage from injuries. Nevertheless, we did not exclude that Inflares might have other functions, e.g., regulating neuronal activities or affecting tissue recovery, that were not directly tested in our system. It is also possible that different densities or patterns of Inflares might drive distinct downstream microglial activities and thus are decoded as multi-strategy responses. This underscores the importance of detecting Inflares as a potential diagnostic marker of the brain internal status to guide further manipulations. Given the tight associations between brain injury and multiple diseases, including diabetes (Hamed, 2017), neurodegenerative diseases (Gardner and Yaffe, 2015) and brain tumors (Chen et al., 2012), and the involvement of microglia in such processes, it is conceivable that Inflares may also be present and contribute to the generation and progression of these diseases. In such situations, targeting pannexin 1 or related upstream/downstream molecules could be of importance for their treatment in the clinical setting.

## Supporting information

Table S1. Statistical analysis

## Acknowledgments

We thank Dr. Yulong Li at Peking University for sharing the ATP1.0 sensor and related plasmid constructs. We thank Dr. Qingchun Guo and his team in the imaging core facility at the Chinese Institute for Brain Research (CIBR) for generous help in imaging experiments and data analysis. We thank Dr. Fei Zhao and his team in the vector core at CIBR for providing related AAV reagents. We thank Dr. Wenzhi Sun at CIBR for help with the two-photon microscope. Some schematics were created with the Biorender website.

## Funding

This work was supported by

The Chinese Brain Initiative Project, National Key Research and Development Program, Project 2021ZD0202200, Subject 2021ZD0202203 (M.J.)

The Project of Intergovernmental Science and Technology Innovative Cooperation, National Key Research and Development Program 2021YFE0116400 (M.J.)

The Beijing Nova Program Z20111000680000 (M.J.)

The Beijing Postdoctoral Science Foundation (J.L.)

## Author contributions

Y.C. performed experiments related to ATP and microglial imaging in FLA injury and ATP signal extraction and characterization. P.L. conducted the cell type screening and MCAO-related experiments. J.L. carried out the microglial morphological analysis and related biochemical experiments. Y.W. performed slice imaging experiments. Z.W. engineered the ATP1.0 sensor and made sensor expressing AAV constructs. All authors contributed to data analysis. Y.C. and M.J. wrote the manuscript with input from all other authors.

## Data and materials availability

The correspondence and material requests should be addressed to M.J.

