## Supplementary materials for "Spatiotemporally selective ATP events from astrocytes encode injury information and guide sustained microglial response"

Materials and Methods

**Animals.** Wild-type C57BL/6J mice (from Charles River or animal facility at CIBR) or Cx3cr1-GFP mice (Cx3cr1^GFP/+^) mice (gift from Dr. Minmin Luo’s lab at CIBR) at 5–7 weeks of age were used for surgery and two-photon imaging. Postnatal day (P)5–7 C57BL/6J mice were used for AAV injection, with subsequent preparation of acute brain slices at 2 weeks after injection. The pannexin 1 conditional knockout mice (panx1^flox/+^) were purchased from GemPharmatech and have a 140-bp deletion in exon 2 of *Panx1*, and homozygous mice were obtained for experiments. The LSL-Cas9-IRES-tdtomato transgenic mice were a gift from Dr. Yulong Li’s lab at Peking University. All mice were either family-housed or pair-housed in a temperature-controlled room with a 12-h light/dark cycle. All procedures for animal surgery and experimentation were performed using protocols approved by the Animal Care & Use Committees of the Chinese Institute for Brain Research (#CIBR-IACUC 007).

**Stereotaxic surgery, virus injection and slice preparation.** Mice were anesthetized with an intraperitoneal (i.p.) injection of avertin (250 mg/kg body weight, Sigma). For in vivo two-photon imaging, a metal head-fixed recording chamber was affixed to the skull of an anesthetized mouse. After a 3- to 4-day recovery, mice were re-anesthetized and placed on a stereotaxic frame (RWD Instruments). A cranial window was opened on the visual cortex with the following coordinates: anterior–posterior, –2.2 mm relative to the bregma; medial–lateral, 2.0 mm relative to the bregma; dorsal–ventral, 0.5 mm below the dura at an angle of 30°. Then 400–500 nl of AAV was injected into the cortex at a slow rate of 30 nl/min with a glass electrode and microsyringe pump (Nanoliter 2000 injector, WPI). After the injection was completed, the glass electrode was left in place for an additional 10 min before it was slowly withdrawn. After the brain surface was cleaned with sterile saline, a 4-mm × 4-mm coverslip was used to replace the skull. Mice were allowed to recover from anesthesia under a heating pad. Two-photon imaging was performed at 3 weeks after AAV injection. The Cx3cr1-GFP mice underwent a similar surgical procedure without AAV injection and were allowed to recover for 1 week before imaging. Neonatal mice at P5–7 were anesthetized by being placed on a pre-chilled metal box and were injected with 300 nl of AAV into the visual cortex; tissue glue was used to suture the skin. Mice were then transferred back to their home cages to allow full recovery from anesthesia.

To prepare acute brain slices, mice were anesthetized 2 weeks after AAV injection and transcardially perfused with 5 ml pre-chilled slicing buffer consisting of (in mM) 110 choline-Cl, 2.5 KCl, 1.25 NaH_2_PO_4_, 25 NaHCO_3_, 7 MgCl_2_, 25 glucose and 2 CaCl_2_. The mice were then decapitated, and their brains were removed immediately and placed in pre-chilled oxygenated slicing buffer (oxygenated with 95% O_2_ + 5% CO_2_). Brains were sectioned into 300-µm slices using a VT1200 vibratome (Leica), and slices containing the visual cortex were transferred immediately to oxygenated ACSF consisting of (in mM) 125 NaCl, 2.5 KCl, 1.25 NaH_2_PO_4_, 25 NaHCO_3_, 1.3 MgCl_2_, 25 glucose and 2 CaCl_2_. Slices were allowed to recover at 33°C for 30 min before being transferred to a chamber for imaging.

Unless stated, all AAVs were made from the vector core of Chinese Institute for Brain Research and aliquoted and stored at −80°C until use. The following AAVs were used in this study: AAV2/9-GfaABC1D-ATP1.0, AAV2/9-GfaABC1D-jRGECO1a, AAV2/9-hSyn-taCaspase3, AAV2/9-hSyn-ATP1.0 (from WZ Bioscience, China), AAV2/9-GfaABC1d-HSV-TK-mCherry, AAV2/9-hSyn-mCherry, AAV2/9-GfaABC1D-hM_3_Dq-mCherry, AAV-hsyn-hM_4_Di-mCherry (from WZ Bioscience), AAV2/9-GfaABC1d-cre-panx1 gRNA, AAV2/9-GfaABC1d-cre-scrambled gRNA and AAV2/9-GfaABC1d-cre.

**Two-photon imaging and focal laser ablation (FLA).** In vivo imaging and acute slice imaging were performed using a two-photon laser-scanning microscope (FVMPE-RS; Olympus) with a pulsed Ti:sapphire laser (Insight DeepSee; Newport Spectra-Physics). Images were acquired by using a 25× water immersion objective (XLPLN25XWMP2, 1.05 NA; Olympus). For imaging ATP dynamics by using the ATP1.0 sensor, the recording frequency was set as 10 Hz and every 10 images were averaged; individual recording periods were typically for 20 min in vivo and 10 min in acute slices. The recording area was typically 500 µm × 500 µm, except when the spatial distribution of Inflares was characterized, during which an extended imaging field of 1400 µm × 1400 µm was recorded and stacked together. To record microglial migration, image stacks were captured with 20-µm thickness using a step size of 5 µm (500 µm × 500 µm × 20 µm).

For FLA, a 920-nm laser (Olympus) was set at its maximum power and used to scan the center of the imaging region (50 µm in diameter) for 2 s. Successful FLA was indicated by the formation of a circle of high fluorescence around the FLA region. For slice experiments, 80% laser power for 1 s was used for FLA. For experiments related to microglial morphological changes, gene expression profiling and related biochemical experiments, an extended FLA was performed in which a region of ~500 µm in diameter was injured to assess the damage at the tissue level.

**Ablation of neurons, microglia and astrocytes.** To ablate neurons, AAV2/9-hSyn-taCaspase3 was injected into the cortex, followed by a 3-week period for expression. The CSF1R inhibitor BLZ945 (MCE, #HY-12768) was added to the food pellets at a concentration of 2 g drug/kg food weight first, and this was fed to the mice twice a day at 3 g pellets per day for 21 consecutive days to effectively remove microglia. The gliotoxin l-α-aminoadipate (L-αAA; Aladdin, A100535) was prepared in ACSF at a stock concentration of 20 mg/ml and injected into the cortex locally in a total volume of 500 nl. The HSV-TK substrate ganciclovir (MCE, #HY-13637A) was injected i.p. at 100 mg/kg of body weight for 14 consecutive days to ablate astrocytes expressing HSV-TK. Successful cell ablation was confirmed by immunostaining of cell markers.

**Drug administration.** Drug administration for in vivo two-photon imaging experiments was as follows. Apyrase (1000 U/ml; A6237-100UN, Sigma,) was prepared in ACSF, and 0.6 µl of the solution was injected into the cortex near the imaging area. The inactivation of apyrase by heat was done at 95°C for 40 minutes. The hM_4_Di synthetic ligand clozapine *N*-oxide (CNO; 250 μM, #A3317, APExBIO) was dissolved in saline and injected i.p. at a volume of 300 µl. The pannexin blocker probenecid (PBN; 4 mg/ml, P36400, Invitrogen) was dissolved in saline and injected i.p. at a concentration of 200 mg/kg body weight 40 min before in vivo imaging or FLA stimulation. For the proliferation experiment, 250 mg/kg of BrdU was injected i.p. 30 min before extended FLA, and the mice were sacrificed 48 h after FLA.

For the acute slice experiments, thapsigargin (TG; 10 μM, HY-13433, MCE) was bath applied for 40 min to deplete astrocytic calcium stores before imaging. CNO (10 µM), PBN (1 mM), trovafloxacin (TROVA; 50 µM, HY-A0170, MCE), ^10^panx1 (100 µM, SML2152, Sigma) and ^sc^panx1 (100 µM, SML2082, Sigma), DCPIB (20 µM, 1540/10, R&D), clodronate (Clo; 10 μM, #HY-107794, MCE) were all bath-applied in the imaging chamber for 5 min before imaging.

**Immunofluorescence staining.** Mice were transcardially perfused with saline, followed by 4% paraformaldehyde (PFA) in phosphate-buffered saline (PBS). Brains were postfixed in 4% PFA at 4°C overnight and were then transferred to 30% sucrose in PBS at 4°C for 48–72 h. Brains were cut on a freezing microtome (CM3050S, Leica), and coronal sections (50 µm) that included the visual cortex were collected. The sections were permeabilized with 0.5% Triton X-100 in PBS, blocked with blocking buffer (5% Bovine albumin (BSA), 0.3% Triton-X100 in PBS) at room temperature for 1 h and incubated with primary antibodies in antibody dilution buffer (1% BSA, 0.3% Triton-X100 in PBS) at 4°C overnight. They were then stained with secondary antibodies at room temperature for 2 h and mounted with 4’,6-Diamidino-2-phenylindole dihydrochloride (DAPI, 0.1 µg/ml) in glycerol/PBS (1:1). For the proliferation experiment, before the immunostaining method described above, an antigen retrieval step for 5-Bromo-2’-deoxyuridine (BrdU) detection was performed with 2 M HCl for 30 min at 37°C, followed by incubation with 0.1 M sodium borate (pH 8.5) for 10 min. Images were obtained with a laser scanning confocal microscope (TCS SP8, Leica).

The following primary antibodies were used in this study (information after antibody name are listed with dilution, catalog # and supplier): rabbit anti-Iba1 (1:1000, ab178846, Abcam), mouse anti-Iba1 (1:500, ab283319, Abcam), chicken anti-GFAP (1:1000, ab4674, Abcam), rabbit anti-GFAP (1:1000, ab7260, Abcam), rabbit anti-NeuN (1:1000, ab177487, Abcam), mouse anti-NeuN (1:1000, ab104224, Abcam), rabbit anti-GFP (1:1000, ab290, Abcam) and rat anti-BrdU (1:1000, ab7260, Abcam,). Secondary antibodies consisted of Alexa Fluor 488–conjugated goat anti–rabbit IgG (1:1000, ab150077, Abcam,), Alexa Fluor 555–conjugated goat anti–rabbit IgG (1:1000, ab150078, Abcam,), Alexa Fluor 488–conjugated goat anti–mouse IgG H&L (1:1000, ab150113, Abcam), Alexa Fluor 555–conjugated goat anti–mouse IgG (1:1000, ab150114, Abcam), Alexa Fluor 647–conjugated goat anti–chicken IgY (1:1000, ab150171, Abcam) and Alexa Fluor 647–conjugated goat anti–rat IgG H&L (1:1000, ab150167, Abcam).

**TUNEL assay.** After FLA in vivo, brain sections were prepared as described above for immunofluorescence staining. Apoptotic cells were detected by terminal deoxynucleotidyl transferase dUTP nick end labeling (TUNEL) assay (Wojciech et al., 1993) using the One Step TUNEL Apoptosis Assay Kit (Beyotime, C1090). Sections were rinsed with PBS and permeabilized with 0.1% Triton X-100 in PBS for 5 min. Slides were rinsed in PBS for 5 min and then incubated at 37°C for 1 h with 50 µl of TUNEL reaction mixture in the dark. Slides were again washed three times with PBS for 10 min and mounted with 0.1 µg/ml DAPI in glycerol/PBS (1:1). Images were obtained with the Leica TCS SP8 and imported into Imaris version 9.7 software for further analysis.

**Western blotting.** Brain cortex tissue with or without the FLA region was isolated from mice treated with saline or PBN. Tissues were homogenized in Radio immunoprecipitation assay (RIPA) lysis buffer (Beyotime, P0013B) to extract total protein, in the presence of a protease inhibitor cocktail (Thermo Scientific, 78429) on ice. Quantification of total protein was conducted using a Bicinchonininc acid (BCA) protein assay (Wiechelman et al., 1988) kit (Biomed, PA101-01). Protein samples were boiled with 5× loading buffer consisting of 300 mM Tris-HCl pH 6.8, 10% Sodium dodecyl sulfate (SDS), 30% glycerol, 0.02% Bromophenol blue at 95°C for 5 min. Approximately 40 µg of protein in each sample was used for SDS polyacrylamide gel-electrophoresis (PAGE), after which the proteins were transferred to polyvinylidene difluoride (PVDF) membranes (Millipore, IPVH00010). The membranes were blocked with 5% BSA in Tris-buffered saline supplemented with 0.1% Tween-20 and incubated with rabbit anti-cleaved caspase-3 (Asp175) (1:1000, #9661, CST), rabbit anti–caspase-3 (1:1000, #9662, CST) and mouse anti–β-actin (1:1000, ab8226, Abcam) primary antibodies at 4°C overnight. Horseradish peroxidase (HRP)-conjugated goat anti–rabbit IgG H&L (1:5000, GAR0072, Multi Science) and HRP-conjugated rabbit anti–mouse IgG H&L (1:1000, ab6728, Abcam) were used at room temperature for 1 h. Signals were detected using the Super ECL Detection Reagent by Chemiluminescent Imaging System (Invitrogen). Quantification of bands was performed using ImageJ (National Institutes of Health).

**Quantitative real-time PCR analysis (qPCR).** Total RNA was extracted from cortex tissues (in the absence and presence of FLA) from wild-type mice treated with saline or PBN or from astrocytic panx1 knockout mice using the RNeasy Mini kit (Qiagen). RNA concentration and purity were evaluated with a NanoDrop One (Thermo Scientific). Reverse transcription was carried out by using the PrimeScript II 1^st^ Strand cDNA Synthesis Kit (Takara). Quantitative PCR was carried out with SYBR Green SuperMix (Transgene) on a CFX96 Touch Real-Time PCR Detection System (Bio-Rad). Relative mRNA expression was calculated using the 2^−ΔΔCT^ method. Each real-time PCR reaction was performed in triplicate. Primers for all genes were designed to cross exon–intron junctions. *Actb* was used as an internal control for samples. Primer sequences are as follows (all 5′–3′, forward and reverse):

*Il1b*, AATGCCACCTTTTGACAGTGAT and GATGTGCTGCTGCGAGATTT;

*Il6*, ACTTCACAAGTCGGAGGCTT and GCCACTCCTTCTGTGACTCC;

*Cxcl10*, AGTGCTGCCGTCATTTTCTG and AAGCTTCCCTATGGCCCTCA;

*Tnf*, CCCTCACACTCACAAACCAC and ATAGCAAATCGGCTGACGGT;

*Ccl2*, ACCTGCTGCTACTCATTCACC and ATTCCTTCTTGGGGTCAGCA;

*Ccl5*, CTCACCATATGGCTCGGACA and GCACTTGCTGCTGGTGTAGA;

*Actb*, GCGGGCGACGATGCT and TCATCTTTTCACGGTTGGCCT;

*Il10* GGTTGCCAAGCCTTATCGGA and CACCTTGGTCTTGGAGCTTATT;

*Tgfb1,* ACGTCACTGGAGTTGTACGG and TTGGGGCTGATCCCGTTGAT;

*Ccl22*, GACCTCTGATGCAGGTCCCTAT and TTGCGGCAGGATTTTGAGGT;

*Ntf3*, TGGTTACTTCTGCCACGATCTTAC and CTTTGATCCATGCTGTTGCCT;

*Bdnf*, GGCTGGAATAGACTCTTGGCA and TCTCACCTGGTGGAACTTCTTTG;

*Ngf*, GAGCGCATCGAGTGACTT and GACCACAGGCCAAAACTCCA;

*Gdnf*, CCGGACGGGACTCTAAGATG and CAGGCATATTGGCGGCG.

**RNA library construction and sequencing.** RNA integrity was assessed using the RNA Nano 6000 Assay Kit of the Bioanalyzer 2100 system (Agilent Technologies, CA, USA). A total amount of 1 µg RNA per sample was used as input material for RNA library preparation(Li et al., 2018) . Briefly, mRNA was purified from total RNA using poly(T) oligo–attached magnetic beads. Fragmentation was carried out using divalent cations under elevated temperature in First Strand Synthesis Reaction Buffer (5×). First-strand cDNA was synthesized using random hexamer primers and M-MuLV Reverse Transcriptase (RNase H–). Second-strand cDNA synthesis was subsequently performed using DNA Polymerase I and RNase H. Remaining overhangs were converted into blunt ends via exonuclease/polymerase activity. After adenylation of the 3′ ends of DNA fragments, adaptors with a hairpin loop structure were ligated to prepare for hybridization. To select cDNA fragments of preferentially 370–420 bp in length, the library fragments were purified with the AMPure XP system (Beckman Coulter, Beverly, USA). Then PCR was performed with Phusion High-Fidelity DNA polymerase, Universal PCR primers and Index (X) Primer. Finally, PCR products were purified (AMPure XP system) and library quality was assessed on the Agilent Bioanalyzer 2100 system. The clustering of the index-coded samples was performed on a cBot Cluster Generation System using TruSeq PE Cluster Kit v3-cBot-HS (Illumina). After cluster generation, the prepared libraries were sequenced on an Illumina NovaSeq platform and 150-bp paired-end reads were generated.

**MCAO in mice.** Adult female C57BL/6J mice weighing 20–25 g were used. Briefly, the common carotid artery, internal carotid artery and external carotid artery were separated. A 6-0 nylon suture coated with silica gel was inserted from the external carotid artery into the internal carotid artery and then was gently inserted into the middle cerebral artery. The success of occlusion was determined by monitoring the decrease in surface cerebral blood flow (CBF) to 30–40% of baseline CBF using a laser speckle contrast imaging system (Simopto, Wuhan, China). Reperfusion was performed by withdrawing the suture 1 h after occlusion. In ATP1.0 imaging experiments, the PBN was injected i.p. into each mouse at two time points, 6 h before MCAO surgery and 6 h after reperfusion, at a concentration of 200 mg/kg body weight. In the behavioral analysis, PBN was injected for three consecutive days at the same concentration.

TTC staining was used to confirm the brain damage after MCAO surgery. The mice were anesthetized and heart-perfused with saline, and their brains were rapidly removed and continuously sectioned into six coronal slices (thickness, 2 mm). Slices were stained with 2% 2,3,5-triphenyltetrazolium chloride (TTC; Sigma, T8877) for 10 min at 37°C in a drying box and were then fixed with 4% PFA for 5 min. Images of the stained sections were captured using a digital camera. The infarction area on each TTC-stained section was measured using Adobe Photoshop CC2018 software (Adobe, USA). Infarction rate (%) = (the area of viable brain tissue in the right hemisphere − the area of viable brain tissue in the left hemisphere) × 100/total area of the slice.

**Behavioral analysis.**	The neurological score of each group was evaluated at 72 h after reperfusion using the longa neural scoring method (Enrique et al., 1989). The scoring system was as follows: 0, normal walk or no neurological deficit; 1, failure to extend opposite forepaw fully or a mild focal neurological deficit; 2, circling to the contralateral side or a moderate focal neurological deficit; 3, falling to the contralateral side or a severe focal neurological deficit; and 4, no spontaneous walking with depressed consciousness.

An open field test was used to detect the motor ability of MCAO model mice treated with saline or PBN for three consecutive days after reperfusion. Mice were put into a chamber (40 cm × 40 cm × 40 cm) and allowed to move freely for 300 s. At the same time, the motion trajectory of each mouse was recorded with an infrared camera recording system (RWD), and motion trajectory, distances and mean speed in zone were analyzed with a SMART video tracking system (Panlab). The balance beam test was used to investigate the balancing ability of MCAO mice treated with saline or PBN for three consecutive days after reperfusion. Mice were put on the balance beam (40 cm × 2 cm), and time spent on the balance beam was recorded (maximum recording time: 120 s).

**Data analysis and statistics.** All in vivo and acute slice imaging data were processed in AQuA60 (Wang et al., 2019). Inflares were further confirmed manually based on processed imaging data. Using the spots function in Imaris software (Oxford Instruments, Imaris 9.7, x64), the number of neurons, microglia, astrocytes and apoptotic cells was determined. To quantify microglial morphological transformations, the length of processes, terminal points of branches, intersections of distance from the soma center and soma volume were measured by using the filament and surface function in Imaris 9.7. Graphs were generated using Origin 95 (Origin Lab software). Animals were randomly assigned to treatment groups. All behavioral experiments and analyses were carried out by investigators who were blind with respect to treatment assignments. All confocal image analyses were performed by a single blinded investigator.

All summary data are reported as the mean ± s.e.m., except where indicated otherwise. Pearson's omnibus normality test was used to evaluate the normality and equal variances between groups. For paired groups, the statistical difference was analyzed using a paired Student’s t-test. For unpaired groups, the F-test was first performed to compare the variances, and, subsequently, a Student’s t-test was conducted. To analyze differences in gene expression after FLA, the Mann-Whitney test was used. Significant differences were set at *p* < 0.05. All detail statistical information is summarized in Supplementary Table 1.

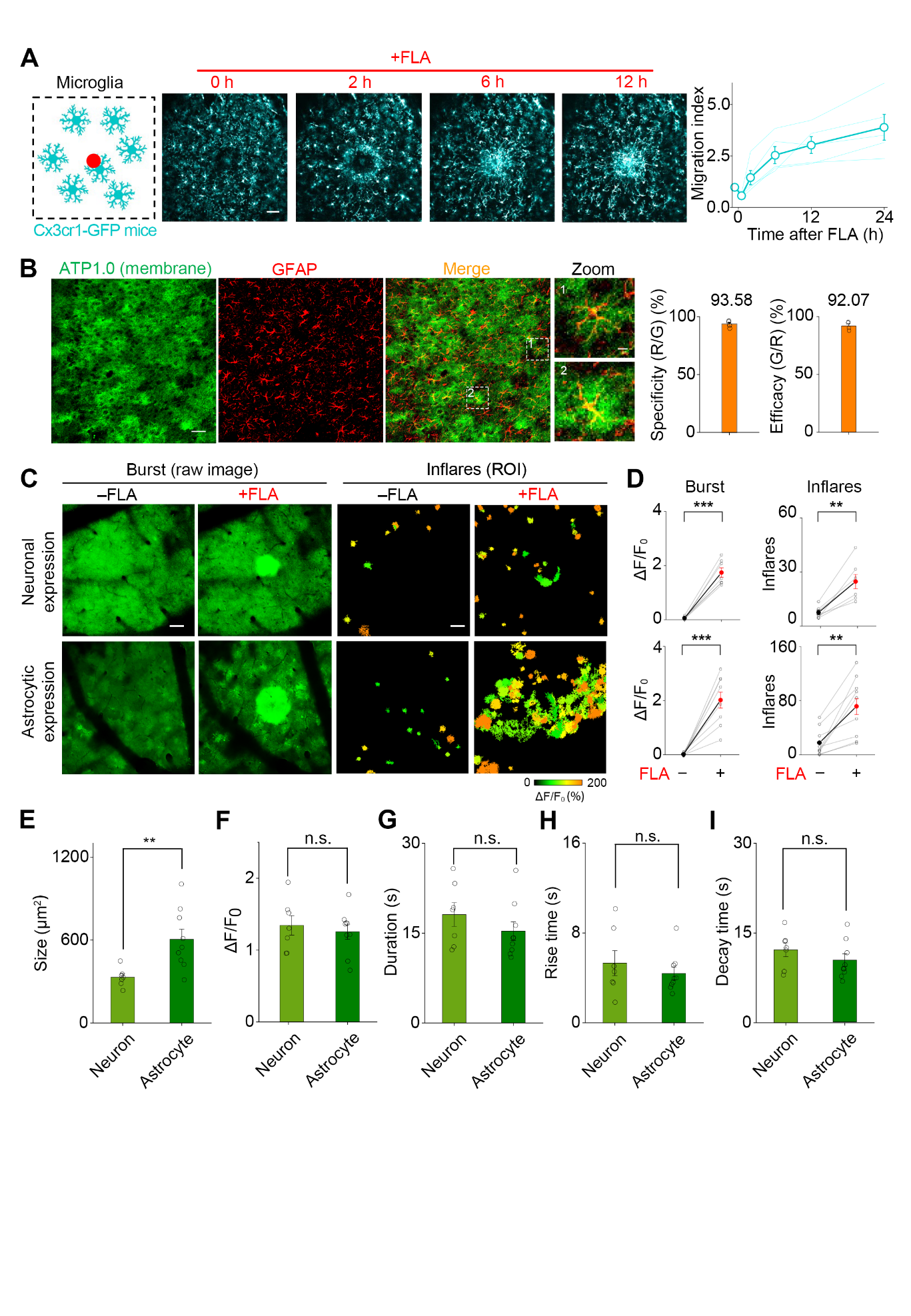

Figure S1. Microglial motility and ATP dynamics after FLA injury.

(A) Two-photon imaging of microglial migration in Cx3cr1-GFP mice after FLA. The migration index was quantified as the relative fluorescence change in the injury center (red circle) over the whole image (*n* = 5 mice).

(B) Astrocyte-specific expression of ATP1.0 as indicated by the colocalization of green fluorescence with GFAP staining (red; *n* = 4 mice). Data are plotted with the colocalization ratio of red (R) over green (G) signal (specificity), or G over R (efficacy).

(C) Representative images of sensor expression and ATP dynamics when expressed in neurons or astrocytes.

(D) Group analysis of the ATP burst (ΔF/F_0_) and the number of Inflares before and after FLA (*n* = 7 and 9 mice for neuronal and astrocytic expression, respectively).

(E-I) Group analysis of inflare properties when ATP1.0 was expressed in neurons or astrocytes (*n* = 7 and 9 mice for neuronal and astrocytic expression, respectively).

Scale bars, 50 µm, except 10 µm in zoomed images in (B). Data are shown as the mean ± s.e.m. ***p* < 0.01; ****p* < 0.001; n.s., not significant. See Table S1 for statistics.

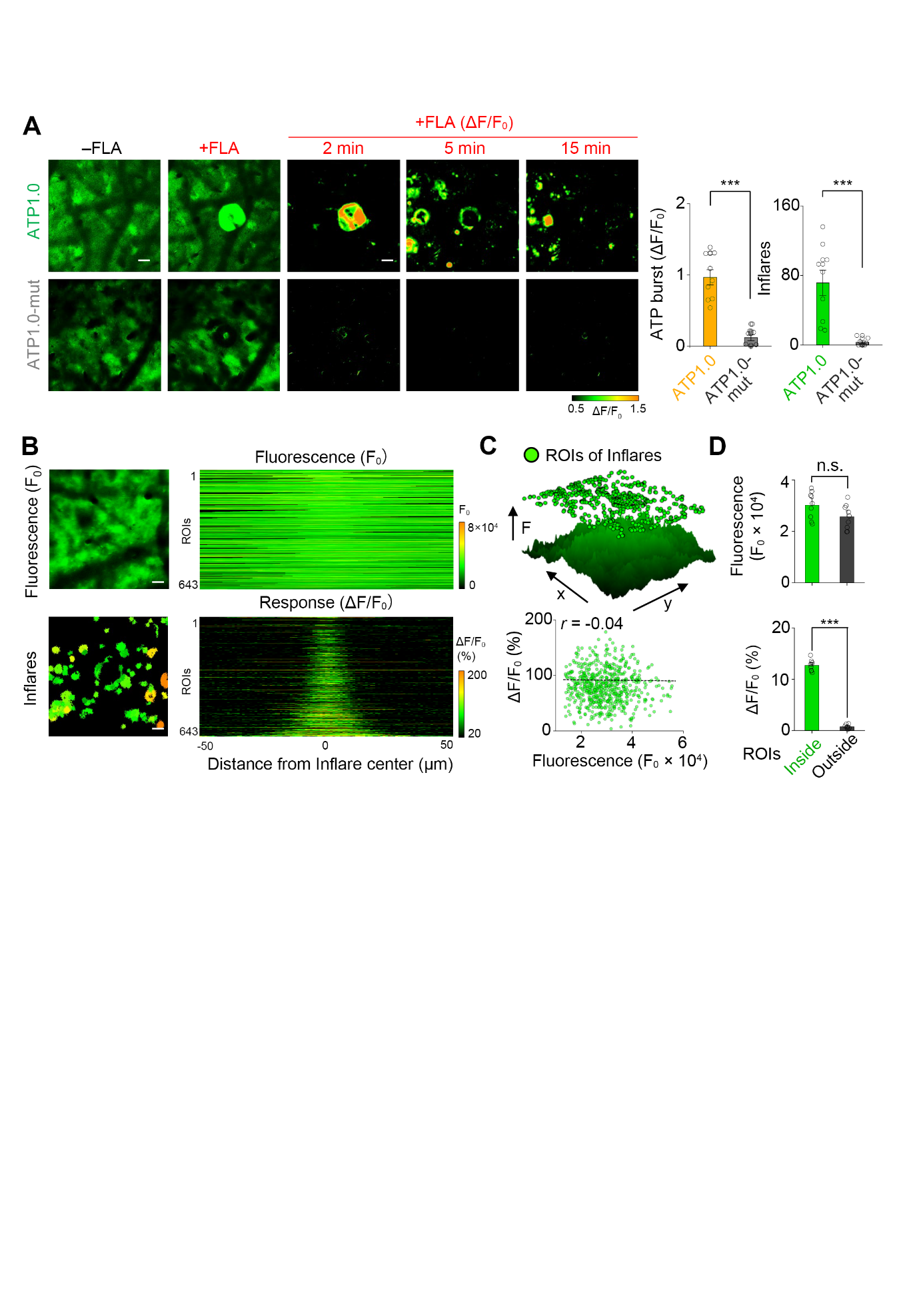

Figure S2. The signal specificity of the fluorescence response after FLA injury.

(A) Representative images and quantification of the sensor expression and ATP dynamics in mice expressing ATP1.0 or ATP1.0-mut in astrocytes (*n* = 9 and 8 mice for ATP1.0 and ATP1.0-mut, respectively).

(B) Left, representative images of basal fluorescence of ATP1.0 (upper) and the ROIs of Inflares after FLA (lower). Right, the spatial distribution of the basal fluorescence (upper) and the fluorescence response (lower). The ROIs were aligned with their center at position 0, and signals were plotted across 100 µm in space (*n* = 643 ROIs from *n* = 9 mice).

(C) The spatial distribution (upper) and the correlation (lower) of the fluorescence responses with corresponding basal fluorescence (F_0_) in ROIs of Inflares (*n* = 643 ROIs from *n* = 9 mice).

(D) A comparison of basal fluorescence (upper) and fluorescence response (ΔF/F_0_, lower) inside and outside of the inflare ROIs (*n* = 9 mice).

Scale bars, 50 µm. Data are shown as the mean ± s.e.m. ****p* < 0.001; n.s., not significant. See Table S1 for statistics.

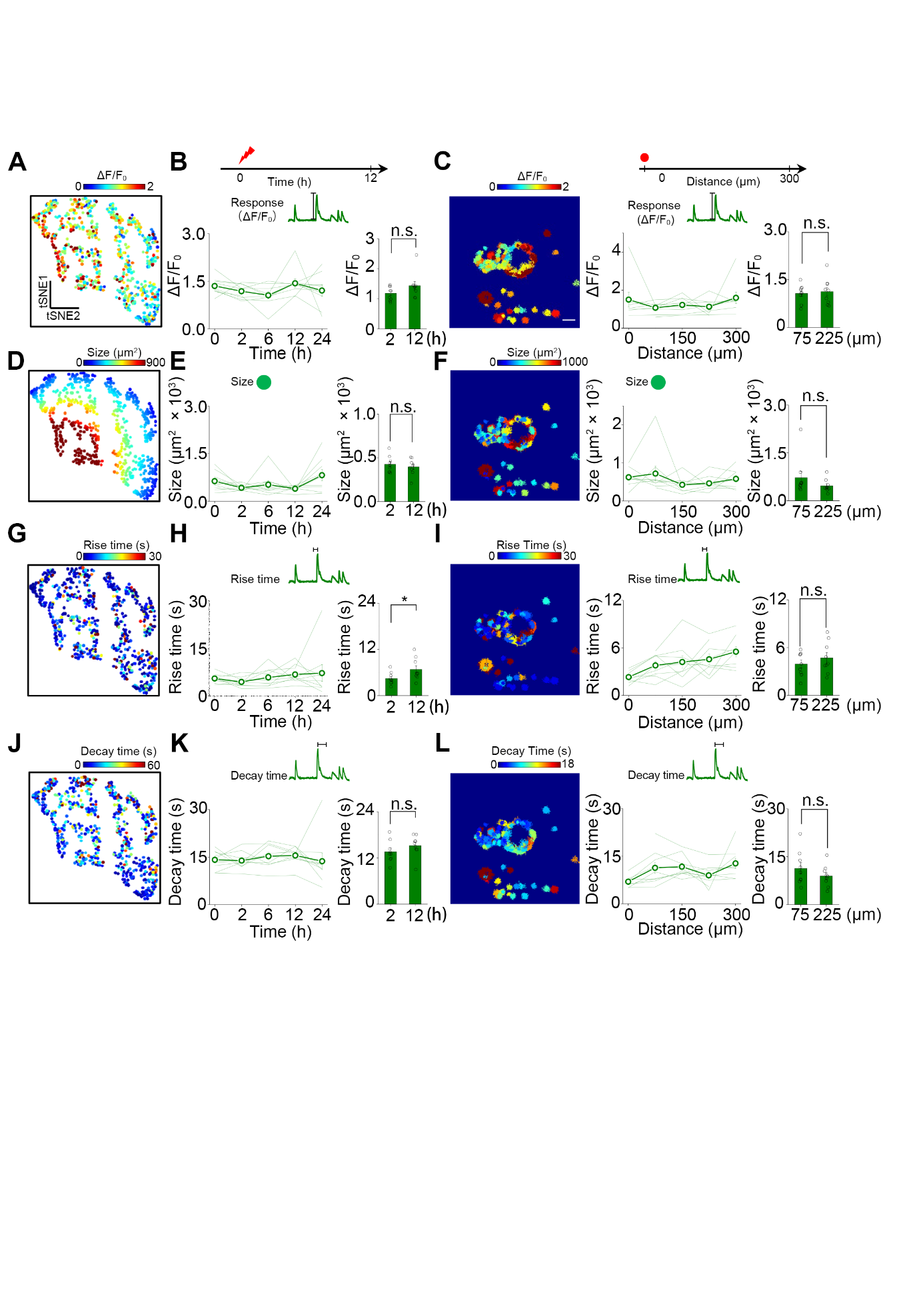

Figure S3. The homogeneous signal properties of Inflares across time and space.

(A) A t-t-distributed stochastic neighbor embedding (tSNE) plot of Inflares according to their peak response. Note that there were no obvious sub-groups within Inflares.

(B, C) The fluorescence response of Inflares at different times after the injury (B) and at different distances from the injury (C) (*n =* 5 mice in b, *n =* 9 mice in c).

(D-F) Similar as (A-C), except inflare size is shown (*n =* 5 mice in e, *n =* 9 mice in f).

(g-i) Similar as (A-C), except inflare rise time is shown (*n =* 5 mice inh, *n =* 9 mice in i).

(j-l) Similar as (A-C), except inflare decay time is shown (*n =* 5 mice ink, *n =* 9 mice in l). Scale bars, 50 µm.

Data are shown as the mean ± s.e.m. **p* < 0.05; n.s., not significant. See Table S1 for statistics.

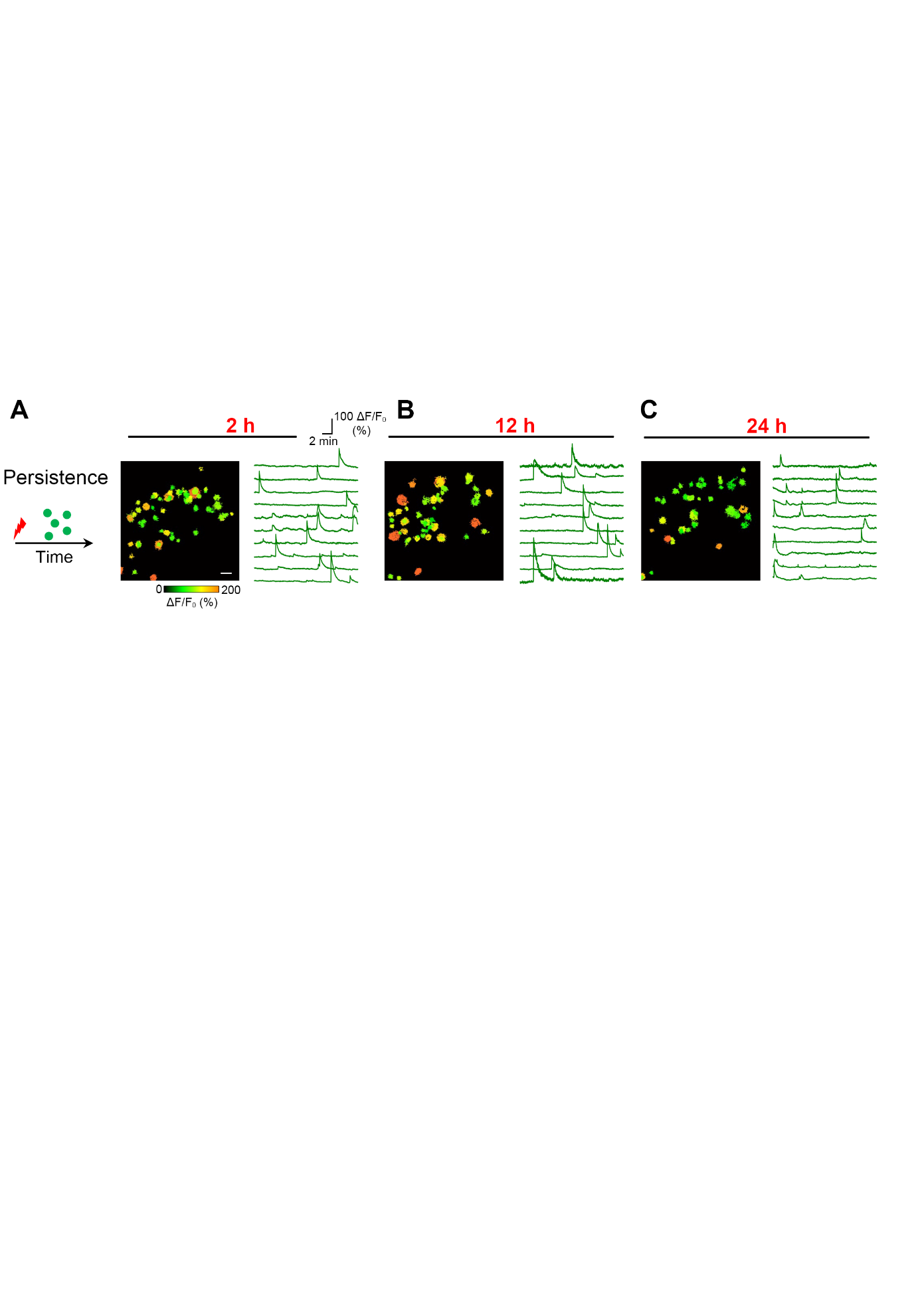

Figure S4. The ROIs and traces of Inflares recorded at the indicated times after injury.

(A-C) The ROIs of Inflares (left) and representative traces (right) at 2 h (A), 12 h (B) and 24 h (C) after FLA injury. Scale bar, 50 µm.

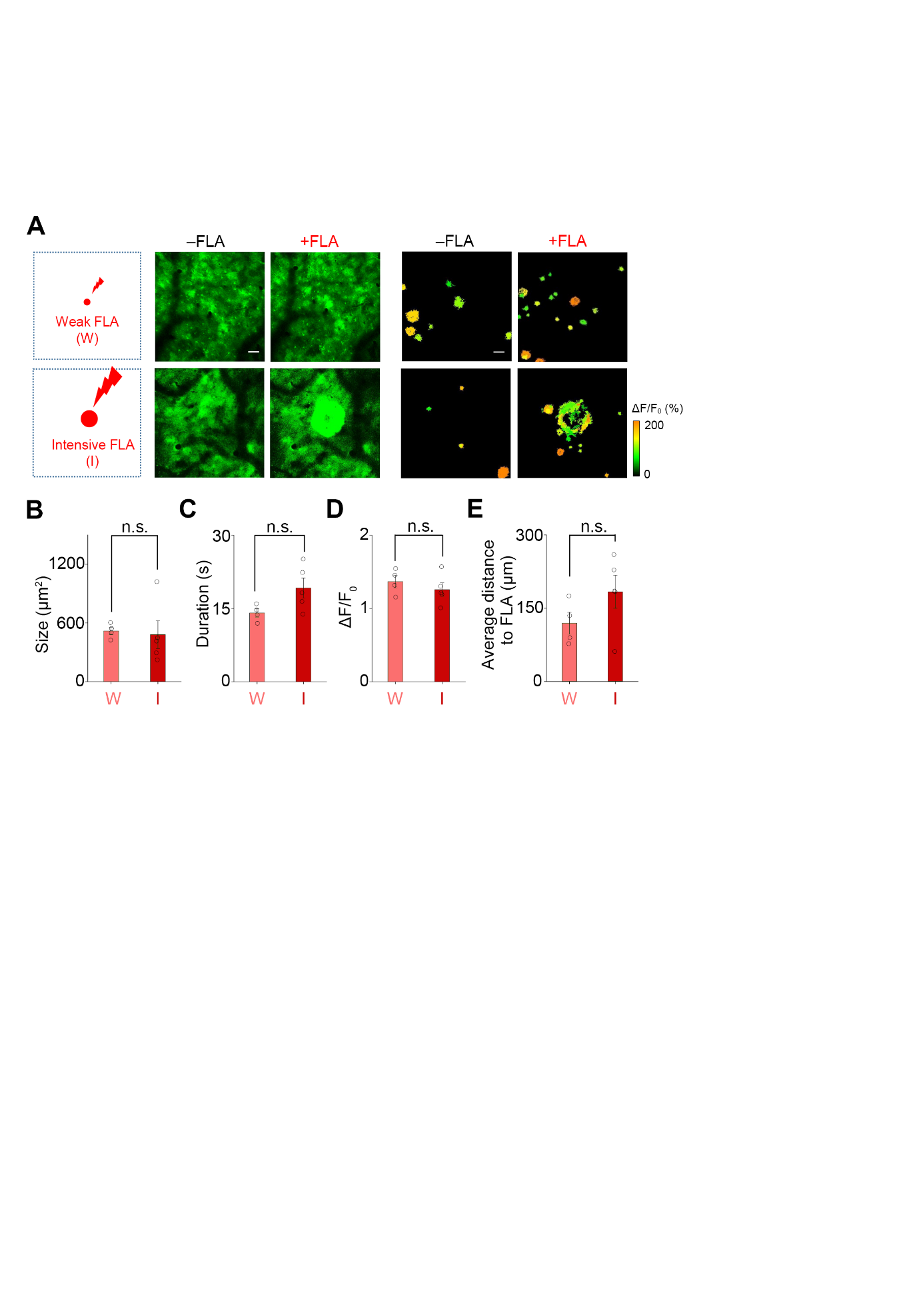

Figure S5. Individual Inflares evoked by different intensities of injury share similar properties.

(A) Schematic (left) of representative fluorescence images (middle) and ROIs of Inflares (right) in ATP1.0-expressing mice in response to weak or intensive FLA injury. The weak and intensive injuries were generated by setting the laser power to 60% for 0.02 s or 100% for 4 s, respectively.

(B-E) Group analysis of inflare properties including signal size (B), duration (C), ΔF/F_0_ (D) and average distance to FLA (E) after weak and intensive FLA (*n* = 5 and 4 mice, respectively).

Scale bars, 50 µm. Data are shown as the mean ± s.e.m. n.s., not significant. See Table S1 for statistics.

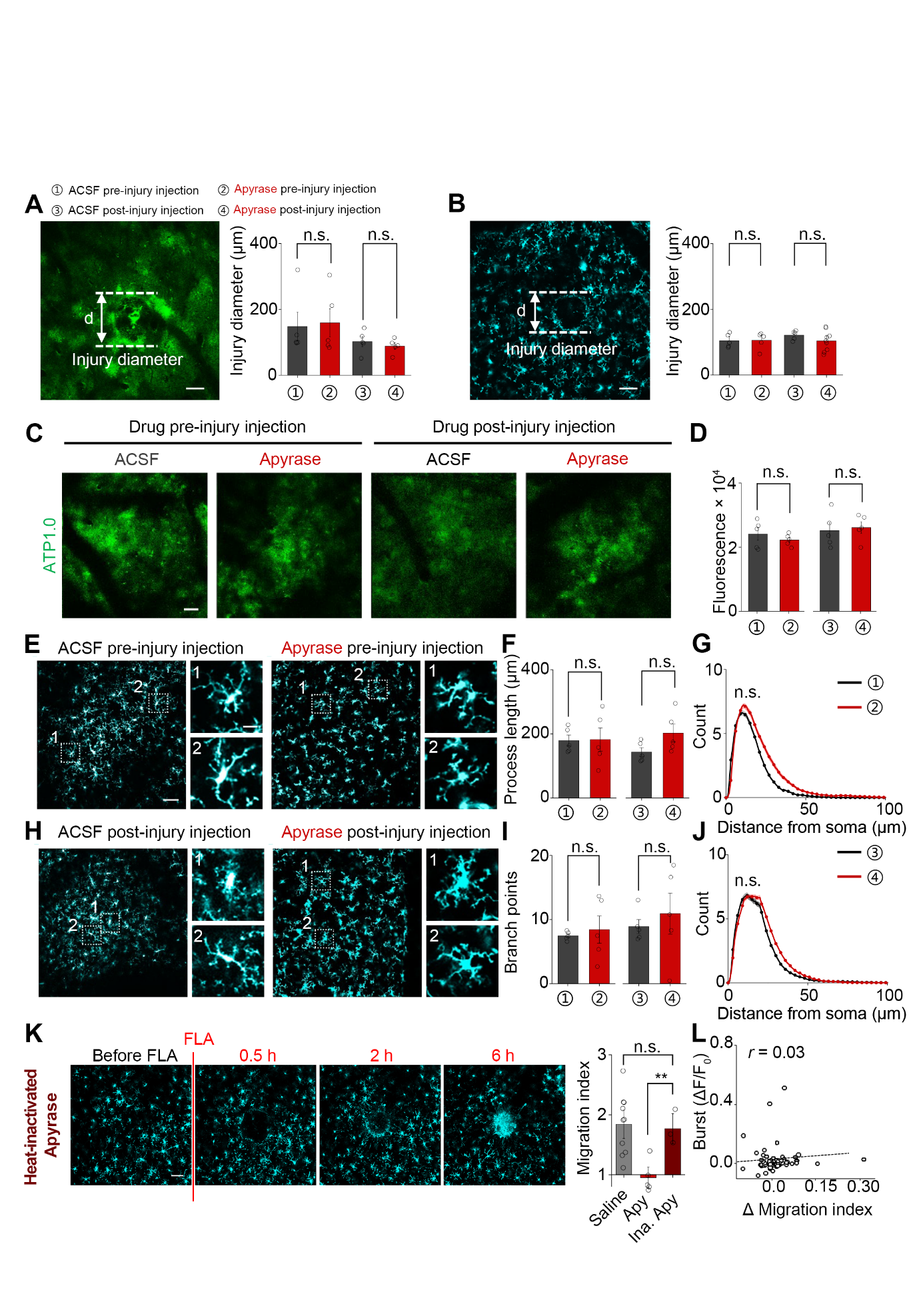

Figure S6. The specificity of ATP blockage on microglial migration and Inflares.

(A, B) Representative images showing the injury size in ATP1.0-expressing mice (A) and Cx3cr1-GFP mice (B). Injury boundaries are indicated by dashed lines, and their diameters (d) were quantified (*n* = 5 mice for ATP1.0 and *n* = 5–9 mice for Cx3cr1-GFP). Different experimental groups are indicated by numbers.

(C, D) Representative images (C) and group analysis (D) of the expression level of ATP1.0 sensors (*n* = 5 mice in each group).

(E–J) Representative images and quantification of microglial morphology under different treatments (*n* = 5 mice in each group).

(K) Microglial migration at the indicated time points in the presence of heat-inactivated apyrase (Ina. Apy; *n* = 10, 5, 3 in Saline, Apy, Ina. Apy, respectively).

(L) The correlation between changes in microglial migration and the fluorescence response to ATP burst (*n* = 5 mice).

Scale bars, 50 µm. Data are shown as the mean ± s.e.m. ***p* < 0.01; n.s., not significant. See Table S1 for statistics.

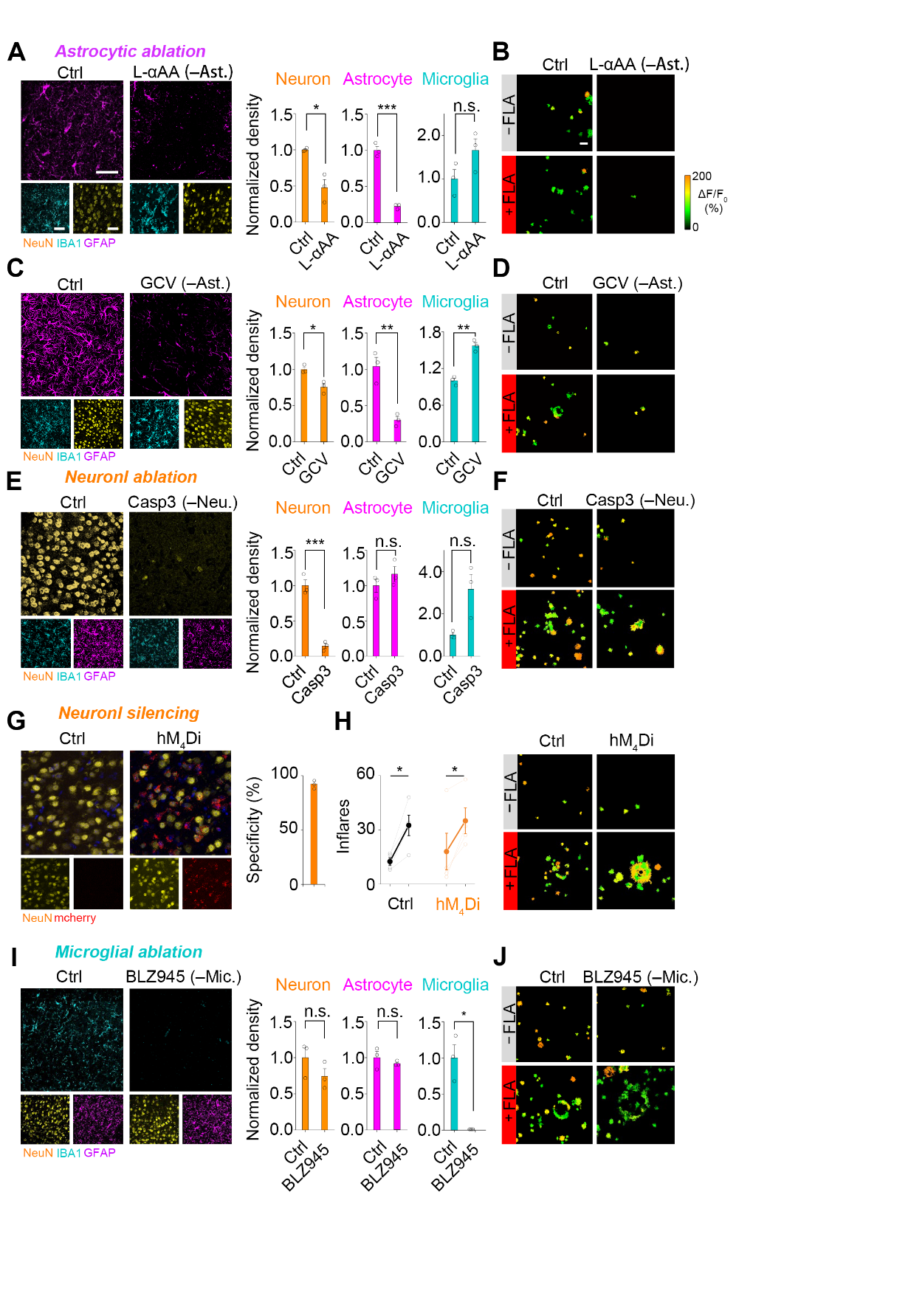

Figure S7. The cellular source of Inflares.

(A) Representative images (left) and group analysis (right) of the selective astrocyte ablation by local injection of the gliotoxin l-αAA near the imaging region. Mice injected with same volume of ACSF were used as the control (*n* = 3 mice in each group).

(B) ROIs of Inflares in control and astrocyte-ablated mice after FLA.

(C, D) Similar to (A and B), except astrocytes were selectively ablated by astrocytic expression of HSV-TK followed by application of GCV. Mice that did not receive GCV but did express HSV-TK were used as the control (*n* = 3 mice in each group).

(E, F) Similar to (a,b), except local neurons were selectively ablated by neuronal expression of ta-Caspase 3. Neuronal expression of mCherry was used as the control (*n* = 3 mice in each group).

(G) Representative images of mice expressing chemogenetic hM_4_Di-mCherry (hM_4_Di) or not (Ctrl). The colocalization of mCherry with a neuronal marker (NeuN) was quantified to determine specificity (right; *n* = 3 mice).

(H) Total inflare number and representative ROIs of Inflares in Ctrl and hM_4_Di-expressing mice after i.p. delivery of CNO (*n* = 4 mice in each group).

(I, J) Similar to (A and B), except the microglia were selectively ablated by oral administration of BLZ945 added to the food pellets. Control mice were fed normal food (*n* = 3 mice in each group).

Scale bars, 50 µm. Data are shown as the mean ± s.e.m. **p* < 0.05; ***p* < 0.01; ****p* < 0.001; n.s., not significant. See Table S1 for statistics.

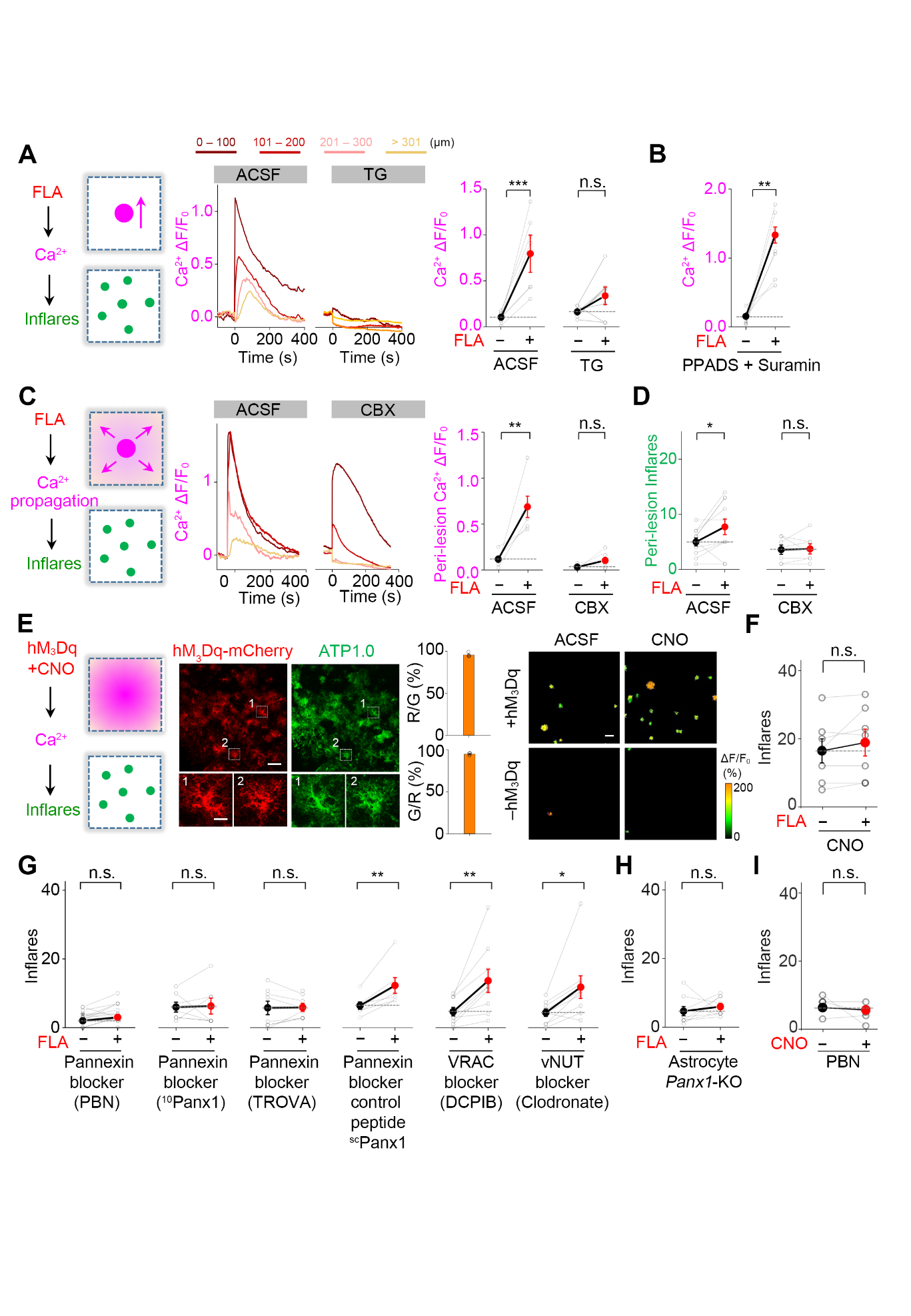

Figure S8. The molecular mechanism responsible for Inflares.

(A) The fluorescence response of astrocytes expressing jRGECO1a at different distances from the FLA. ROIs at different distances from the injury center were selected, with changes in fluorescence plotted over time. Brain slices were bathed in ACSF or thapsigargin (TG) (*n* = 9 slices from 3 mice (9/3) for ACSF and *n* = 7/3 for TG). Traces with different colors represent the fluorescence response quantified from ROIs within the indicated distance to injury (e.g., 0-100 µm from injury center).

(B) The fluorescence response of jRGECO1a after FLA in slices treated with broad P2 receptor blockers (PPADS + suramin) (*n* = 9 slices from 3 mice). (c) Similar to (a), except slices were bathed in ACSF or the gap junction blocker carbenoxolone (CBX), and signals located 100 µm away from the injury (referred to as the peri-lesion region) were quantified (*n* = 6 slices from 2 mice (6/2) for ACSF and *n* = 6/3 for CBX).

(D) Group analysis of inflare number from the peri-lesion region in slices treated with ACSF or CBX (*n* = 11 slices from 2 mice (11/2) for ACSF and *n* = 7/3 for CBX).

(E) From left to right: Schematic illustration of experiments; Representative images and quantification of the co-expression of hM_3_Dq-mCherry and ATP1.0 in astrocytes; ROIs of of Inflares in hM_3_Dq-expressing or control (mCherry alone, −hM_3_Dq) slices in response to CNO application.

(F) The inflare number in CNO-treated slices in the absence and presence of FLA injury (with hM_3_Dq expression; *n* = 7 slices from 3 mice).

(G) Inflares in response to different pharmacological treatments (*n* = 13 slices from 4 mice (13/4) for PBN, *n* = 7/3 for ^10^panx1, *n* = 8/3 slices/mice for TROVA, *n* = 7/3 for ^sc^panx1, *n* = 9/3 for DCPIB, *n* = 9/3 for clodronate).

(H) Number of Inflares when pannexin 1 was selectively knocked out (KO) in astrocytes (*n* = 9 slices from 2 mice for astrocyte Panx1-KO). The panx1^flox^ mice were injected with AAV-GFaABC1d-cre to selectively knock out Panx1.

(I) Inflare number in PBN-treated slices in the presence of CNO (with hM_3_Dq expression; *n* = 4 slices from 2 mice).

Scale bars, 50 µm. Data are shown as the mean ± s.e.m. **p* < 0.05; ***p* < 0.01; ****p* < 0.001; n.s., not significant. See Table S1 for statistics.

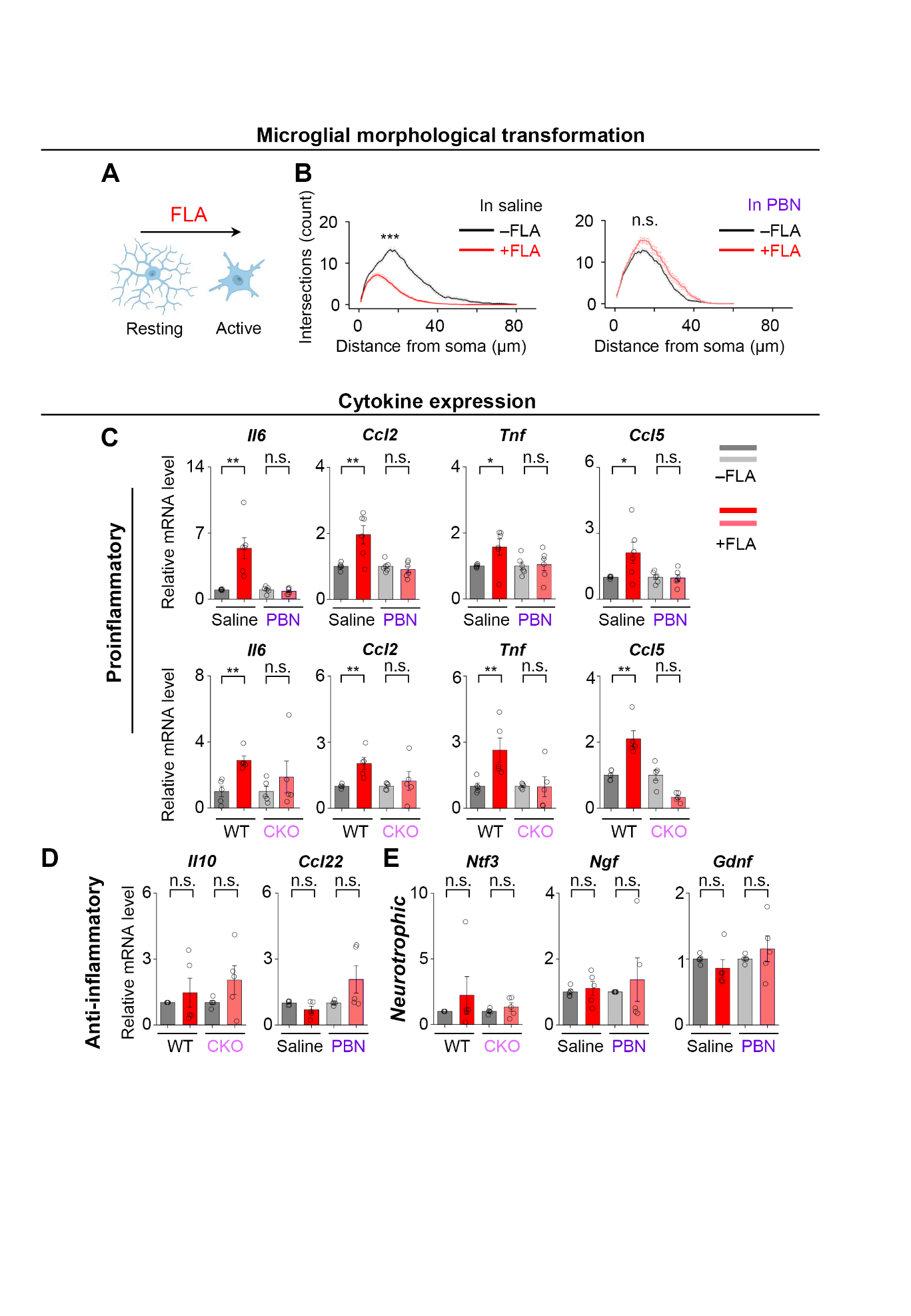

Figure S9. Injury causes broad activity changes in microglia in an inflare-dependent manner.

(A) Cartoon showing the typical microglial morphology at rest and after activation following FLA.

(B) Branch intersections at different distances from the soma (Sholl analysis) were quantified (saline: *n* = 60/5 cells/mice for −FLA, *n* = 73/5 for +FLA; PBN: *n* = 43/5 for −FLA, *n* = 41/5 for +FLA).

(C) Expression of genes encoding pro-inflammatory cytokines and chemokines in saline- and PBN-treated mice (*n* = 6 mice in each group; upper) and in wild-type (WT) and astrocytic panx1 knockout (CKO) mice (*n* = 5 mice in each group) at 24 h after FLA.

(D, E) Similar to (C), except genes encoding anti-inflammatory (D) and neurotrophic factors (E) were quantified (*n* = 5 mice in each group).

Scale bars, 50 µm. Data are shown as the mean ± s.e.m. **p* < 0.05; ***p* < 0.01; ****p* < 0.001; n.s., not significant. See Table S1 for statistics.

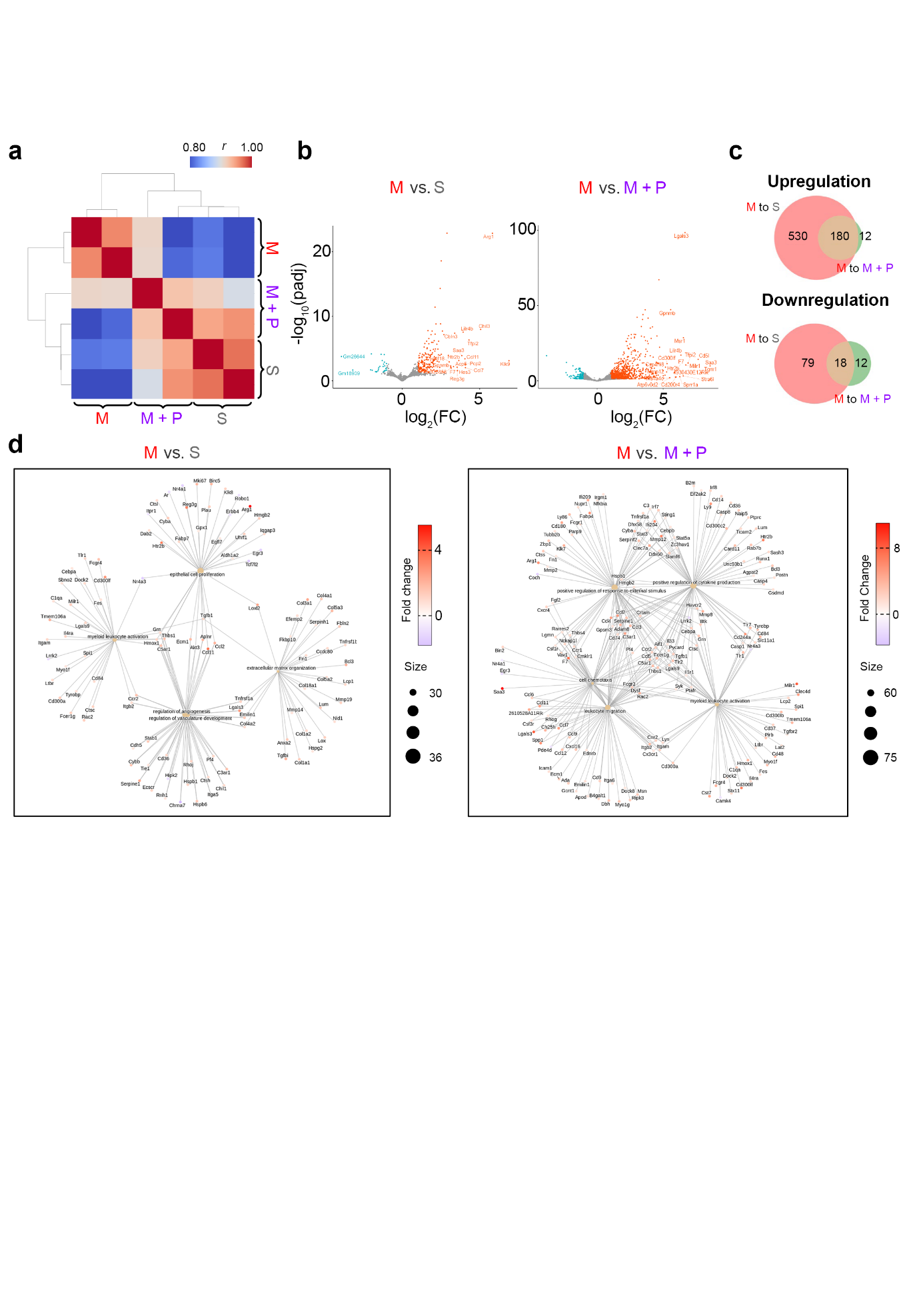

Figure S10. RNA sequencing of mouse brains subjected to MCAO.

(A) A correlation analysis of RNA sequencing results from different groups (two biological replicates in each group).

(B) Volcano plot of changes in gene expression between MCAO and sham mice (left) and between MCAO and MCAO + PBN mice (right). Genes with a significant increase or decrease in each group are labeled in red or blue, respectively.

(C) Venn diagram showing the number of genes that were up-regulated (upper) or down-regulated (lower) when comparing MCAO vs. sham or MCAO vs. MCAO + PBN mice.

(D) The network analysis of genes showing significant changes between MCAO vs. sham mice (left) and MCAO vs. MCAO + PBN mice (right).
Video S1. ATP dynamics before and after FLA recorded by ATP1.0 fluorescent sensor in vivo

**Video S2. Intensive Inflares recorded at 12 h after ischemic stroke in mice**
