## Supplementary material for "Spatiotemporally selective ATP events from astrocytes encode injury information and guide sustained microglial response": Table S1. Statistical analysis

Supplementary Table 1 Statistical analysis

| Figure | Sample size (from left to right) | Statistical test | P values | F and DF |
| --- | --- | --- | --- | --- |
| 1E | n = 9 mice (B) n = 9 mice (I) | Unpaired two-tailed t-test | p = 0.0014 (B vs I) | F = 8.3 × 10^-41^; DF = 8 |
| 1F | n = 9 mice (B) n = 9 mice (I) | Unpaired two-tailed t-test | p = 6.4 × 10^-6^ (B vs I) | F = 5.2 × 10^-06^; DF = 8 |
| 1G | n = 9 mice (B) n = 9 mice (I) | Unpaired two-tailed t-test | p = 0.29 (B vs I) | F = 1.5 ×10^-07^; DF = 8 |
| 1H | n = 9 mice (B) n = 9 mice (I) | Unpaired two-tailed t-test | p = 4.1 × 10^-6^ (B vs I) | F = 1.2 ×10^-15^; DF = 8 |
| 1I | n = 9 mice (±FLA, ATP1.0) n = 8 mice (±FLA, ATP1.0-Mut) n = 9 mice (±FLA, ATP1.0) n = 8 mice (±FLA, ATP1.0-Mut) | Paired two-tailed t-test | p = 1.5 × 10^-4^ (±FLA, ATP1.0, Burst) p = 0.05 (±FLA, ATP1.0-Mut, Burst) p = 0.0021 (±FLA, ATP1.0, Inflares) p = 0.16 (±FLA, ATP1.0-Mut, Inflares) | DF = 8 DF = 7 DF = 8 DF = 7 |
| 2B | n = 9 mice (B) n = 9 mice (I) | Unpaired two-tailed t-test | p = 2.7 × 10-6 (B vs I) | F = 1.0 × 10^-13^; DF = 3 |
| 2D | n = 5 mice (B) n = 5 mice (I) | Unpaired two-tailed t-test | p = 2.7553 x 10^-6^ (B vs I) | F = 0.0087;  DF = 4 |
| 2F (Duration of directionality) | n = 9 mice (B) n = 9 mice (I) | Unpaired two-tailed t-test | p = 0.0038 (B vs I) | F = 3.7 × 10^-5^; DF = 8.45 |
| 2F (Region of directionality) | n = 9 mice (B) n = 9 mice (I) | Unpaired two-tailed t-test | p = 0.0018 (B vs I) | F = 6.1 × 10^-16^; DF = 8.0 |
| 2H | n = 5 mice (–FLA) n = 5 mice (+FLA) n = 5 mice (Random | Paired two-tailed t-test | p = 0.028 (–FLA vs +FLA) p = 1.4 × 10^-4^ ^(^+FLA vs Random) p = 0.058 (–FLA vs Random) | DF = 4 DF = 4 DF = 4 |
| 2I | n = 5 mice (I) n = 4 mice (W) | Unpaired two-tailed t-test | p = 0.031 | F = 0.25; DF = 7 |
| 2J | n = 27 mice | Spearman's rank correlation coefficient | p = 0.0069 |  |
| 3J | n = 10 mice (A, Burst) n = 5 mice (B, Burst) n = 5 mice (C, Burst) n = 10 mice (A, Inflares) n = 5 mice (B, Inflares) n = 5 mice (C, Inflares) n = 10 mice (A, Migration) n = 5 mice (B, Migration) n = 9 mice (C, Migration) | Unpaired two-tailed t-test | p = 0.0050 (A vs B, Burst) p = 0.67 (A vs C, Burst) p = 4.0 × 10^-8^ (A vs B, inflares) p = 1.6 × 10^-6^ (A vs C, inflares) p = 0.0028 (A vs B, Migration) p = 0.0039 (A vs C, Migration) | F = 0.44; DF = 13 F = 0.44; DF = 13 F = 0.53; DF = 13 F = 0.51; DF = 13 F = 0.23; DF = 13 F = 0.30; DF = 17 |
| 3K | n = 10 ATP recording  n = 10 Microglia recording | Pearson correlation | p = 5.5 × 10^-6^ (Raw) p = 0.60 (Shuffled) | *r =* 0.75 (Raw) *r =* 0.10 (Shuffled) |
| 4C | n = 6 mice (±FLA, Ctrl) n = 6 mice (±FLA, GCV) n = 7 mice (±FLA, L-αAA) | Paired two-tailed t-test | p = 0.0050 (±FLA, Ctrl)  p = 0.028 (±FLA, GCV)  p = 0.56 (±FLA, L-αAA) | DF = 5  DF = 5  DF = 6 |
| 4E | n = 5 mice (±FLA, Ctrl) n = 4 mice (±FLA, Casp3) | Paired two-tailed t-test | p = 0.045 (±FLA, Ctrl) p = 0.011 (±FLA, Casp3) | DF = 4 DF = 3 |
| 4G | n = 8 mice (±FLA, Ctrl) n = 9 mice (±FLA, BLZ945) | Paired two-tailed t-test | p = 0.011 (±FLA, Ctrl) p = 0.0019 (±FLA, BLZ945) | DF = 7 DF = 8 |
| 4I | n = 7 slices from 3 mice (±FLA, ACSF) n = 7 slices from 3 mice (±FLA, TG) | Paired two-tailed t-test | p = 0.0049 (±FLA, ACSF) p = 0.086 (±FLA, TG) | DF = 6 DF = 6 |
| 4J | n = 7 slices from 3 mice (±CNO, +hM_3_Dq) n = 8 slices from 2 mice (±CNO, -hM_3_Dq) | Paired two-tailed t-test | p = 0.0041 (±CNO, +hM_3_Dq) p = 34 (±CNO, -hM_3_Dq) | DF = 6 DF = 7 |
| 4K | n = 7 slices from 3 mice (ACSF) n = 13 slices from 4 mice (PBN) n = 7 slices from 3 mice (^10^PanX1) n = 8 slices from 3 mice (TROVA) n = 7 slices from 3 mice (^SC^PanX1) n = 9 slices from 3 mice (DCPIB) n = 9 slices from 3 mice (Clodronate) | Unpaired two-tailed t-test | p = 2.0 × 10^-4^ (ACSF vs PBN) p = 0.0098 (ACSF vs 10PanX1) p = 7.6 × 10^-4^ (ACSF vs TROVA) p = 0.36 (ACSF vs SCPanX1) p = 0.69 (ACSF vs DCPIB) p = 0.82 (ACSF vs Clodronate) | F = 0.21; DF = 18 F = 0.43; DF = 12 F = 0.57; DF = 13 F = 0.93; DF = 12 F = 0.10; DF = 14 F = 0.12; DF = 14 |
| 4L | n = 4 mice (±FLA, *Panx1* gRNA) n = 6 mice (±FLA, Scrambled gRNA) | Paired two-tailed t-test | p = 0.39 (±FLA, PanX1 gRNA) p = 0.004 (±FLA, Scrambled gRNA) | DF = 3 DF = 5 |
| 5A | n = 5 mice (Saline) n = 10 mice (PBN) | Unpaired two-tailed t-test | p = 9.8 × 10^-7^ (Saline vs PBN) | F = 0.74; DF = 13 |
| 5B (Branch points) | n = 63 cells from 6 mice (–FLA, Saline) n = 63 cells from 6 mice (+FLA, Saline) n = 52 cells from 5 mice (–FLA, PBN) n = 55 cells from 6 mice (+FLA, PBN) | Unpaired two-tailed t-test | p = 2.1 × 10^-26^ (+FLA, Saline vs PBN) | F = 2.9 × 10^-17^;  DF = 63 |
| 5B (Process length) | n = 63 cells from 6 mice (–FLA, Saline) n = 63 cells from 6 mice (+FLA, Saline) n = 52 cells from 5 mice (–FLA, PBN) n = 55 cells from 6 mice (–FLA, PBN) | Unpaired two-tailed t-test | p = 2.9 × 10^-24^ (+FLA, Saline vs PBN) | F = 3.1 × 10^-15^;  DF =64 |
| 5B (Cell volume) | n = 61 cells from 6 mice (–FLA, Saline) n = 61 cells from 6 mice (+FLA, Saline) n = 61 cells from 6 mice (–FLA, PBN) n = 61 cells from 6 mice (–FLA, PBN) | Unpaired two-tailed t-test | p = 8.0 × 10^-6^ (+FLA, Saline vs PBN) | F = 1.2 × 10^-10^;  DF =80 |
| 5C (Microglia proliferation cell number) | n = 5 mice (Saline) n = 5 mice (PBN) | Unpaired two-tailed t-test | p=0.0043 (Saline vs PBN) | F=0.0017 DF = 4.09 |
| 5C (Proliferation ratio) | n = 5 mice (Saline) n = 5 mice (PBN) | Unpaired two-tailed t-test | p=0.0024 (Saline vs PBN) | F=6.9 × 10^-4^; DF = 4.14 |
| 5D | n = 4 mice (–FLA, Saline) n = 6 mice (+FLA, Saline) n = 5 mice (–FLA, PBN) n = 4 mice (+FLA, PBN) | Unpaired two-tailed t-test | p = 0.013 (±FLA, Saline) p = 0.10 (±FLA, PBN) p = 0.018 (+FLA, Saline vs PBN) | F = 0; DF=5 F = 0.0016; DF = 3.078 F = 0.0032; DF = 5.15 |
| 5E (*Il1b*) | n = 6 mice (–FLA, Saline) n = 6 mice (+FLA, Saline) n = 6 mice (–FLA, PBN) n = 6 mice (+FLA, PBN) n = 5 mice (–FLA, Wt) n = 5 mice (+FLA, Wt) n = 5 mice (–FLA, CKO) n = 5 mice (+FLA, CKO) | Wilcoxon rank sum test | p = 0.0025 (±FLA, Saline) p = 0.345 (±FLA, PBN) p = 0.0061 (±FLA, Wt) p = 0.105 (±FLA, CKO) |  |
| 5E (*Cxcl10*) | n = 6 mice (+FLA, Saline) n = 6 mice (+FLA, Saline) n = 6 mice (+FLA, PBN) n = 6 mice (+FLA, PBN) n = 5 mice (+FLA, Wt) n = 5 mice (+FLA, Wt) n = 5 mice (+FLA, CKO) n = 5 mice (+FLA, CKO) | Wilcoxon rank sum test | p = 0.0025 (±FLA, Saline) p = 0.189 (±FLA, PBN) p = 0.0468 (±FLA, Wt) p = 0.105 (±FLA, CKO) |  |
| 5F (*Il10*) | n = 5 mice (+FLA, Saline) n = 5 mice (+FLA, Saline) n = 5 mice (+FLA, PBN) n = 5 mice (+FLA, PBN) | Wilcoxon rank sum test | p = 0.0061 (±FLA, Saline) p = 0.0694 (±FLA, PBN) |  |
| 5F (*Tgfb1*) | n = 5 (–FLA, Saline) n = 5 (+FLA, Saline) n = 5 (–FLA, PBN) n = 5 (–FLA, PBN) | Wilcoxon rank sum test | p=0.011 (±FLA, Saline) p=0.0060 (±FLA, PBN) |  |
| 5F (*Ntf3*) | n = 5 mice (+FLA, Saline) n = 5 mice (+FLA, Saline) n = 5 mice (+FLA, PBN) n = 5 mice (+FLA, PBN) n = 5 mice (+FLA, Wt) n = 5 mice (+FLA, Wt) n = 5 mice (+FLA, CKO) n = 5 mice (+FLA, CKO) | Wilcoxon rank sum test | p = 0.0061 (±FLA, Saline) p = 0.338 (±FLA, PBN) p = 0.328 (±FLA, Wt) p = 0.265 (±FLA, CKO) |  |
| 5F (*Bdnf*) | n = 5 (–FLA, Saline) n = 5 (+FLA, Saline) n = 5 (–FLA, PBN) n = 5 (–FLA, PBN) | Wilcoxon rank sum test | p=0.0061 (±FLA, Saline) p=0.34 (±FLA, PBN) |  |
| 5G (TUNEL^+^ cell number) | n = 6 mice (Saline, –FLA) n = 6 mice (Saline, +FLA) n = 6 mice (PBN, –FLA) n = 6 mice (PBN, +FLA) | Unpaired two-tailed t-test | p = 0.0060 (+FLA, Saline vs PBN) | F =2.9 × 10-4;  DF = 5 |
| 5G (Cleaved caspase3) | n = 6 mice (Saline, +FLA) n = 6 mice (PBN, +FLA) n = 6 mice (KO, +FLA) | Unpaired two-tailed t-test | p = 0.0229 (Saline vs PBN) p = 0.0288 (Saline vs KO) | F = 0.3066; DF = 10 F = 0.1308; DF = 8 |
| 6A (Blood flow change) | n = 5 mice (Sham)  n = 9 mice (MCAO) | Unpaired two-tailed t-test | p = 1.0 × 10^-8^ (Sham vs MCAO) | F = 0.043; DF = 11 |
| 6A (Infarction rate) | n = 6 mice (Sham)  n = 6 mice (MCAO) | Unpaired two-tailed t-test | p = 6.0 × 10^-5^ (Sham vs MCAO) | F = 0.012;  DF = 6 |
| 6D | n = 6 mice (Sham) n = 5 mice (MCAO) n = 4 mice (PBN) n = 4 mice (Mut) | Unpaired two-tailed t-test | p = 0.72 (±Surgery, Sham) p = 0.0024 (±Surgery, MCAO) p = 0.30 (±Surgery, PBN) p = 0.85 (±Surgery, mut) | F = 0.45; DF = 10 F =1.5 × 10^-4^; DF = 4 F = 0.086; DF = 6 F = 0.65; DF = 6 |
| 6F | n = 10 mice (Sham) n = 9 mice (MCAO) n = 12 mice (MCAO+PBN) | Unpaired two-tailed t-test | p = 0.0062 (MCAO vs PBN) | F = 0.58;  DF = 19 |
| 6G | n = 10 mice (Sham) n = 9 mice (MCAO) n = 12 mice (MCAO+PBN) | Unpaired two-tailed t-test | p = 8.1 × 10^-4^ (MCAO vs PBN) | F = 0.10;  DF = 19 |
| 6H | n = 10 mice (Sham) n = 9 mice (MCAO) n = 12 mice (MCAO+PBN) | Unpaired two-tailed t-test | p = 0.027 (MCAO vs PBN) | F = 0.39;  DF = 18 |
| 6I | n = 10 mice (Sham) n = 9 mice (MCAO) n = 12 mice (MCAO+PBN) | Unpaired two-tailed t-test | p = 0.0063 (MCAO vs PBN) | F = 1.6 × 10^-4^;  DF = 11 |
| Figure S 1D (Burst) | n = 7 mice (Neuron expression) n = 9 mice (Astrocyte expression) | Paired two-tailed t-test | p = 5.80 × 10^-5^ (±FLA, Neuron) p = 1.53 × 10^-4^ (±FLA, Astrocyte) | DF = 6 DF = 8 |
| Figure S1D (Inflares) | n = 7 mice (Neuron expression) n = 9 mice (Astrocyte expression) | Paired two-tailed t-test | p = 0.0017 (±FLA, neuron) p = 0.0021 (±FLA, Astrocyte) | DF = 6 DF = 8 |
| Figure S1E | n = 7 mice (Neuron) n = 9 mice (Astrocyte) | Unpaired two-tailed t-test | p = 0.0055 (Neuron vs Astrocyte) | F = 0.0087;  DF = 10 |
| Figure S1F | n = 7 mice (Neuron) n = 9 mice (Astrocyte) | Unpaired two-tailed t-test | p = 0.61 (Neuron vs Astrocyte) | F = 0.72;  DF = 14 |
| Figure S1G | n = 7 mice (Neuron) n = 9 mice (Astrocyte) | Unpaired two-tailed t-test | p = 0.28 (Neuron vs Astrocyte) | F = 0.75;  DF = 14 |
| Figure S1H | n = 7 mice (Neuron) n = 9 mice (Astrocyte) | Unpaired two-tailed t-test | p = 0.44 (Neuron vs Astrocyte) | F = 0.17;  DF = 14 |
| Figure S1I | n = 7 mice (Neuron) n = 9 mice (Astrocyte) | Unpaired two-tailed t-test | p = 0.26 (Neuron vs Astrocyte) | F = 0.98;  DF = 14 |
| Figure S2A  (Burst) | n = 9 mice (ATP1.0) n = 8 mice (ATP1.0-Mut) | Unpaired two-tailed t-test | p = 8.7 × 10^-6^ (ATP1.0 vs ATP1.0-mut) | F = 0.033;  DF = 11 |
| Figure S2A  (Inflares) | n = 9 mice (ATP1.0) n = 8 mice (ATP1.0-Mut) | Unpaired two-tailed t-test | p = 0.0016 (ATP1.0 vs ATP1.0-mut) | F = 6.1 × 10^-6^;  DF = 8 |
| Figure S2C | n = 643 ROIs from n = 9 mice | Pearson correlation | p = 0.26 | r = -0.04 |
| Figure S2D (Fluorescence) | n = 9 mice | Unpaired two-tailed t-test | p = 0.072 (Inside vs outside) | F = 0.80;  DF = 16 |
| Figure S2D  (ΔF/F_0_) | n = 9 mice | Unpaired two-tailed t-test | p = 2.6 × 10^-11^ (Inside vs outside) | F = 0.0077;  DF = 10 |
| Figure S3B | n = 9 mice (2 h) n = 9 mice (12 h) | Unpaired two-tailed t-test | p = 0.13 (2 h vs 12 h) | F = 0.11;  DF = 16 |
| Figure S3C | n = 9 mice (75 µm) n = 9 mice (225 µm) | Unpaired two-tailed t-test | p = 0.74 (75 µm vs 225 µm) | F = 0.31;  DF = 16 |
| Figure S3E | n = 9 mice (2 h) n = 9 mice (12 h) | Unpaired two-tailed t-test | p = 0.52 (2 h vs 12 h) | F = 0.92;  DF = 16 |
| Figure S3F | n = 9 mice (75 µm) n = 9 mice (225 µm) | Unpaired two-tailed t-test | p = 0.25 (75 µm vs 225 µm) | F = 0.0062;  DF = 9 |
| Figure S3H | n = 9 mice (2 h) n = 9 mice (12 h) | Unpaired two-tailed t-test | p = 0.044 (2 h vs 12 h) | F = 0.011;  DF = 16 |
| Figure S3I | n = 9 mice (75 µm) n = 9 mice (225 µm) | Unpaired two-tailed t-test | p = 0.35 (75 µm vs 225 µm) | F = 0.43;  DF = 16 |
| Figure S3K | n = 9 mice (2 h) n = 9 mice (12 h) | Unpaired two-tailed t-test | p = 0.23 (2 h vs 12 h) | F = 0.97;  DF = 16 |
| Figure S3L | n = 9 mice (75 µm) n = 9 mice (225 µm) | Unpaired two-tailed t-test | p = 0.27 (75 µm vs 225 µm) | F = 0.21;  DF = 16 |
| Figure S5B | n = 5 mice (±FLA, W) n = 4 mice (±FLA, I) | Unpaired two-tailed t-test | p = 0.81 (W vs I, Size) | F = 0.21; DF = 16 |
| Figure S5C | n = 5 mice (±FLA, W) n = 4 mice (±FLA, I) | Unpaired two-tailed t-test | p = 0.073 (W vs I, Duration) | F = 0.13; DF = 7 |
| Figure S5D | n = 5 mice (±FLA, W) n = 4 mice (±FLA, I) | Unpaired two-tailed t-test | p = 0.42 (W vs I, ΔF/F0) | F = 0.79; DF = 7 |
| Figure S5E | n = 5 mice (±FLA, W) n = 4 mice (±FLA, I) | Unpaired two-tailed t-test | p = 0.18 (W vs I, Averaged distance) | F = 0.42; DF = 7 |
| Figure S6A | n = 5 mice (①) n = 5 mice (②) n = 5 mice (③) n = 5 mice (④) | Unpaired two-tailed t-test | p = 0.86 (① vs ②)  p = 0.47 (③ vs ④) | F = 0.99; DF = 8  F = 0.43; DF = 8 |
| Figure S6B | n = 5 mice (①) n = 5 mice (②) n = 5 mice (③) n = 9 mice (④) | Unpaired two-tailed t-test | p = 0.93 (① vs ②)  p = 0.26 (③ vs ④) | F = 0.68; DF = 8  F = 0.15; DF = 12 |
| Figure S6D | n = 5 mice (①) n = 5 mice (②) n = 5 mice (③) n = 5 mice (④) | Unpaired two-tailed t-test | p = 0.40 (① vs ②)  p = 0.76 (③ vs ④) | F = 0.13; DF = 8 F = 0.58; DF = 8 |
| Figure S6F | n = 5 mice (①) n = 5 mice (②) n = 5 mice (③) n = 5 mice (④) | Unpaired two-tailed t-test | p = 0.94 (① vs ②)  p = 0.093 (③ vs ④) | F = 0.18; DF = 8  F = 0.20; DF = 8 |
| Figure S6G | n = 308 cells from 5 mice (①) n = 236 cells from 5 mice (②) | Kolmogorov–Smirnov test | p = 0.20 (① vs ②) |  |
| Figure S6I | n = 5 mice (①) n = 5 mice (②) n = 5 mice (③) n = 5 mice (④) | Unpaired two-tailed t-test | p = 0.68 (① vs ②)  p = 0.57 (③ vs ④) | F = 0.0023; DF = 4 F = 0.061; DF = 8 |
| Figure S6J | n = 689 cells from 5 mice (③)  n = 368 cells from 5 mice (④) | Kolmogorov–Smirnov test | p = 0.60 (③ vs ④) |  |
| Figure S6K | n = 10 mice (Saline) n = 5 mice (Apyrase) n = 3 mice (Heat-inactivated apyrase) | Unpaired two-tailed t-test | p = 0.81 (Saline vs Ina. Apy.) p = 0.0065 (Apyrase vs Ina. Apy.) | F=0.55; DF=6 F=0.78; DF=11 |
| Figure S6L | n = 10 ATP recording  n = 10 Microglia recording | Pearson correlation | p = 0.76 | r = 0.034 |
| Figure S7A | n = 3 mice (Ctrl, Neuron) n = 3 mice (L-αAA, Neuron) n = 3 mice (Ctrl, Astrocyte) n = 3 mice (L-αAA, Astrocyte) n = 3 mice (Ctrl, Microglia) n = 3 mice (L-αAA, Microglia) | Unpaired two-tailed t-test | p = 0.041 (Ctrl vs L-αAA, Neuron)  p = 1.8 × 10^-4^ (Ctrl vs L-αAA, Astrocyte) p = 0.13 (Ctrl vs L-αAA, Microglia) | F = 0.029; DF = 2 F = 0.25; DF = 4 F = 0.80; DF = 4 |
| Figure S7C | n = 3 mice (Ctrl, Neuron) n = 3 mice (GCV, Neuron) n = 3 mice (Ctrl, Astrocyte) n = 3 mice (GCV, Astrocyte) n = 3 mice (Ctrl, Microglia) n = 3 mice (GCV, Microglia) | Unpaired two-tailed t-test | p = 0.013 (Ctrl vs GCV, Neuron)  p = 0.0051 (Ctrl vs GCV, Astrocyte) p = 0.0011 (Ctrl vs GCV, Microglia) | F = 0.64; DF = 4 F = 0.26; DF = 4 F = 0.80; DF = 4 |
| Figure S7E | n = 3 mice (Ctrl, Neuron) n = 3 mice (Casp3, Neuron) n = 3 mice (Ctrl, Astrocyte) n = 3 mice (Casp3, Astrocyte) n = 3 mice (Ctrl, Microglia) n = 3 mice (Casp3, Microglia) | Unpaired two-tailed t-test | p = 6.8 × 10^-4^ (Ctrl vs Casp3, Neuron)  p = 0.31 (Ctrl vs Casp3, Astrocyte) p = 0.089 (Ctrl vs Casp3, Microglia) | F = 0.34; DF = 4 F = 0.86; DF = 4 F = 0.043; DF = 2 |
| Figure S7H | n = 4 mice (±FLA, Ctrl) n = 4 mice (±FLA, hM4Di) | Paired two-tailed t-test | p = 0.029 (±FLA, Ctrl)  p = 0.038 (±FLA, hM4Di) | DF = 3 DF = 3 |
| Figure S7I | n = 3 mice (Ctrl, Neuron) n = 3 mice (BLZ945, Neuron) n = 3 mice (Ctrl, Astrocyte) n = 3 mice (BLZ945, Astrocyte) n = 3 mice (Ctrl, Microglia) n = 3 mice (BLZ945, Microglia) | Unpaired two-tailed t-test | p = 0.22 (Ctrl vs BLZ945, Neuron)  p = 0.38 (Ctrl vs BLZ945, Astrocyte) p = 0.032 (Ctrl vs BLZ945, Microglia) | F = 0.72; DF = 4 F = 0.096; DF = 4 F = 0; DF = 2 |
| Figure S8A | n = 9 slices from 3 mice (±FLA, ACSF) n = 7 slices from 3 mice (±FLA, TG) | Paired two-tailed t-test | p = 2.7 × 10^-4^ (±FLA, ACSF) p = 0.11 (±FLA, TG) | DF = 8 DF = 6 |
| Figure S8B | n = 9 slices from 3 mice (PPADS + Suramin) | Paired two-tailed t-test | p = 0.0012 (±FLA, PPADS + Suramin) | DF = 8 |
| Figure S8C | n = 6 slices from 3 mice (±FLA, ACSF) n = 6 slices from 3 mice (±FLA, CBX) | Paired two-tailed t-test | p = 0.0083 (±FLA, ACSF) p = 0.14 (±FLA, CBX) | DF = 5 DF = 5 |
| Figure S8D | n = 11 slices from 3 mice (±FLA, ACSF) n = 7 slices from 3 mice (±FLA, CBX) | Paired two-tailed t-test | p = 0.026 (±FLA, ACSF) p = 0.85 (±FLA, CBX) | DF = 10 DF = 6 |
| Figure S8F | n = 7 slices from 3 mice (±FLA, CNO) | Paired two-tailed t-test | p = 0.20 (±FLA, CNO) | DF = 6 |
| Figure S8G | n = 7 slices from 3 mice (ACSF) n = 13 slices from 4 mice (PBN) n = 7 slices from 3 mice (^10^PanX1) n = 8 slices from 3 mice (TROVA) n = 7 slices from 3 mice (^SC^PanX1) n = 9 slices from 3 mice (DCPIB) n = 9 slices from 3 mice (Clodronate) | Paired two-tailed t-test | p = 0.0034 (±FLA, ACSF) p = 0.080 (±FLA, PBN) p = 0.89 (±FLA, 10panx1) p = 0.91 (±FLA, TROVA) p = 0.0049 (±FLA, scPanx1) p = 0.0073 (±FLA, DCPIB) p = 0.021 (±FLA, Clodronate) | DF = 7 DF = 12 DF = 6  DF = 7  DF = 6 DF = 8 DF = 8 |
| Figure S8H | n = 9 slices from 2 mice (±FLA, Astrocyte panx1-ko) | Paired two-tailed t-test | p = 0.46 (±FLA, Astrocyte panx1-ko) | DF = 8 |
| Figure S8I | n = 4 slices from 2 mice (±CNO, PBN) | Paired two-tailed t-test | p = 0.31 (±CNO, PBN) | DF = 3 |
| Figure S9B | n = 60 cells from 5 mice (–FLA, Saline) n = 73 cells from 6 mice (+FLA, Saline) n = 43 cells from 6 mice (–FLA, PBN) n = 41 cells from 6 mice (–FLA, PBN) | Kolmogorov–Smirnov test | p = 7.58 × 10^-8^ (Saline) p = 0.28 (PBN) |  |
| Figure S9C  (*Il6*, PBN) | n = 6 (–FLA, Saline) n = 6 (+FLA, Saline) n = 6 (–FLA, PBN) n = 6 (–FLA, PBN) | Wilcoxon rank sum test | p=0.0025 (±FLA, Saline) p=0.76 (±FLA, PBN) |  |
| Figure S9C  (*Ccl2*, PBN) | n = 6 (–FLA, Saline) n = 6 (+FLA, Saline) n = 6 (–FLA, PBN) n = 6 (–FLA, PBN) | Wilcoxon rank sum test | p=0.023 (±FLA, Saline) p=0.76 (±FLA, PBN) |  |
| Figure S9C  (*Tnf*, PBN) | n = 6 (–FLA, Saline) n = 6 (+FLA, Saline) n = 6 (–FLA, PBN) n = 6 (–FLA, PBN) | Wilcoxon rank sum test | p=0.033 (±FLA, Saline) p=0.34 (±FLA, PBN) |  |
| Figure S9C  (*Ccl5*, PBN) | n = 6 (–FLA, Saline) n = 6 (+FLA, Saline) n = 6 (–FLA, PBN) n = 6 (–FLA, PBN) | Wilcoxon rank sum test | p=0.033 (±FLA, Saline) p=0.66 (±FLA, PBN) |  |
| Figure S9C  (*Il6*, CKO) | n = 5 (–FLA, Wt) n = 5 (+FLA, Wt) n = 5 (–FLA, CKO) n = 5 (–FLA, CKO) | Wilcoxon rank sum test | p=0.0061 (±FLA, Wt) p=0.27 (±FLA, CKO) |  |
| Figure S9C  (*Ccl2*, CKO) | n = 5 (–FLA, Wt) n = 5 (+FLA, Wt) n = 5 (–FLA, CKO) n = 4 (–FLA, CKO) | Wilcoxon rank sum test | p=0.0056 (±FLA, Wt) p=0.34 (±FLA, CKO) |  |
| Figure S9C  (*Tnf*, CKO) | n = 5 (–FLA, Wt) n = 5 (+FLA, Wt) n = 5 (–FLA, CKO) n = 5 (–FLA, CKO) | Wilcoxon rank sum test | p=0.0061 (±FLA, Wt) p=0.73 (±FLA, CKO) |  |
| Figure S9C  (*Ccl5*, CKO) | n = 5 (–FLA, Wt) n = 5 (+FLA, Wt) n = 5 (–FLA, CKO) n = 5 (–FLA, CKO) | Wilcoxon rank sum test | p=0.0061 (±FLA, Wt) p=0.99 (±FLA, CKO) |  |
| Figure S9D  (*Il10*, CKO) | n = 5 mice (+FLA, Wt) n = 5 mice (+FLA, Wt) n = 5 mice (+FLA, CKO) n = 5 mice (+FLA, CKO) | Wilcoxon rank sum test | p = 0.328 (±FLA, Wt) p = 0.102 (±FLA, CKO) |  |
| Figure S9D  (*Ccl22*, PBN) | n = 5 (–FLA, Saline) n = 5 (+FLA, Saline) n = 5 (–FLA, PBN) n = 5 (–FLA, PBN) | Wilcoxon rank sum test | p=0.95 (±FLA, Saline) p=0.0718 (±FLA, PBN) |  |
| Figure S9E  (*Ntf3*, CKO) | n = 5 mice (+FLA, Wt) n = 5 mice (+FLA, Wt) n = 5 mice (+FLA, CKO) n = 5 mice (+FLA, CKO) | Wilcoxon rank sum test | p = 0.328 (±FLA, Wt) p = 0.265 (±FLA, CKO) |  |
| Figure S9E  (*Ngf*, PBN) | n = 5 (–FLA, Saline) n = 5 (+FLA, Saline) n = 5 (–FLA, PBN) n = 5 (–FLA, PBN) | Wilcoxon rank sum test | p=0.42 (±FLA, Saline) p=0.75 (±FLA, PBN) |  |
| Figure S9E  (*Gdnf*, PBN) | n = 5 (–FLA, Saline) n = 5 (+FLA, Saline) n = 5 (–FLA, PBN) n = 5 (–FLA, PBN) | Wilcoxon rank sum test | p=0.072 (±FLA, Saline) p=0.80 (±FLA, PBN) |  |
